# Endothelial mTOR maintains hematopoiesis during aging

**DOI:** 10.1101/2020.03.13.990911

**Authors:** Pradeep Ramalingam, Michael G. Poulos, Michael C. Gutkin, Lizabeth Katsnelson, Ana G. Freire, Elisa Lazzari, Jason M. Butler

## Abstract

Aging leads to a decline in hematopoietic stem and progenitor cell (HSPC) function. We recently discovered that aging of bone marrow endothelial cells (BMECs) leads to an altered crosstalk between the BMEC niche and HSPCs, that instructs young HSPCs to behave as aged HSPCs. Here, we demonstrate aging leads to a decrease in mTOR signaling within BMECs that potentially underlies the age-related impairment of their niche activity. Our findings reveal that pharmacological inhibition of mTOR using Rapamycin has deleterious effects on hematopoiesis. To formally determine whether endothelial-specific inhibition of mTOR can influence hematopoietic aging, we conditionally deleted mTOR in ECs (mTOR***^(ECKO)^***) of young mice and observed that their HSPCs displayed attributes of an aged hematopoietic system. Transcriptional profiling of HSPCs from mTOR***^(ECKO)^*** mice revealed that their transcriptome resembled aged HSPCs. Notably, during serial transplantations, exposure of wild type HSPCs to an mTOR***^(ECKO)^*** microenvironment was sufficient to recapitulate aging-associated phenotypes, confirming the instructive role of EC-derived signals in governing HSPC aging.

**Summary:** Ramalingam et al. demonstrate that pharmacological inhibition of mTOR adversely impacts aging hematopoiesis. The authors demonstrate that aging results in decreased mTOR signaling within the bone marrow endothelium and endothelial-specific inhibition of mTOR causes hematopoietic defects observed during physiological aging.

## Introduction

The number of elderly is increasing with unprecedented speed around the globe. The aging process is associated with an increased susceptibility to cardiovascular and hematopoietic disorders. Aging is associated with increased risk of negative outcomes/treatment failures because elderly patients respond poorly to myeloablative strategies that are necessary for the successful transplantation of hematopoietic stem and progenitor cells (HSPCs) and also develop prolonged cytopenias following myelosuppressive therapies that are often used to treat hematopoietic malignancies and other cancers (Balducci, 2003). One of the most significant changes observed during the aging process is a decline in the overall function of endothelial cells (ECs), including the EC niche of the hematopoietic system (Das et al., 2018; El Assar et al., 2012; Le Couteur and Lakatta, 2010). An increasing body of evidence demonstrating functional interactions between the HSPCs and its niche suggests that both local and systemic factors regulate HSPC function (Bowers et al., 2018; Crane et al., 2017; Decker et al., 2018; Lazzari and Butler, 2018; Pinho and Frenette, 2019). However, to date, most reports describing alterations in the aged hematopoietic compartment have focused on the cell-intrinsic properties of HSPCs. For instance, it has been demonstrated that whereas the absolute number of immunophenotypically defined HSPCs increases with age, aged HSPCs exhibit a decrease in their long-term reconstitution abilities (Chambers et al., 2007; Geiger et al., 2013; Kowalczyk et al., 2015; Pang et al., 2011; Rossi et al., 2005) and show a significant myeloid bias at the expense of lymphopoiesis (Cho et al., 2008; Dykstra and de Haan, 2008; Rossi et al., 2005; Van Zant and Liang, 2003). In contrast, the role of the aged microenvironment – specifically, aged bone marrow ECs (BMECs) – in regulating HSPC function during aging has been far less examined.

It has been shown that BMECs assume an instructive role in supporting HSPC self-renewal and differentiation into lineage-committed progeny, in part mediated by activation of their AKT signaling pathway (Butler et al., 2010; Poulos et al., 2015). When interrogating signaling pathways downstream of AKT, we found that the mechanistic target of Rapamycin (mTOR) signaling pathway stimulated the expression of pro-HSPC paracrine factors within AKT-activated ECs (Kobayashi et al., 2010). The mTOR complex utilizes many signals, including growth factors and oxygen tension, to regulate cell growth, proliferation, protein synthesis, energy metabolism, and survival (Zoncu et al., 2011). mTOR activity is strongly linked to physiological aging, and inhibiting mTOR activity increases the longevity of aged mice (Harrison et al., 2009; Inoki et al., 2003; Kaeberlein et al., 2005; Kapahi et al., 2004; Lee et al., 2010; Vellai et al., 2003; Wullschleger et al., 2006). Physiological aging of the HSPC pool is also regulated by mTOR activity, and it has been reported that the mTOR pathway is dysregulated in aged mice and that increased mTOR activation within HSPCs results in their depletion (Chen et al., 2009b). However, the regulation of HSPC activity by mTOR signaling in the aged BM endothelial niche and its contribution to the aging of the hematopoietic system has not been studied.

We have recently demonstrated that aged BMECs can instruct young HSPCs to function as aged HSPCs, whereas young BMECs can preserve the functional output of aged HSPCs (Poulos et al., 2017). Upon further examination of the pathways regulating BMEC niche function, we found that physiological aging is associated with decreased AKT/mTOR signaling within BMECs, which potentially impairs their niche activity. In support of this hypothesis, we observed that pharmacological inhibition of mTOR signaling in aged mice by Rapamycin treatment resulted in an increase in hematopoietic aging phenotypes at homeostasis and severe defects in the hematopoietic system following myelosuppression. Furthermore, EC-specific deletion of mTOR in young mice resulted in premature aging of the hematopoietic system, where many of the phenotypic and functional attributes of HSPCs from mTOR***^(ECKO)^*** mice closely resembled aged mice. Together, these studies have identified BMECs and AKT/mTOR signaling as key components in regulating HSPC self-renewal and differentiation during the aging process.

## Results and Discussion

To determine whether physiological aging alters mTOR signaling within the bone marrow (BM), we performed immunoblot analysis in whole bone marrow (WBM) cells which revealed that aging is associated with a decreased expression of mTOR signaling targets including phospho-S6 and phospho-4EBP-1 **(Supplemental Figure 1A-B)**. Phosphoflow analysis demonstrated a decreased expression of phospho-mTOR, phospho-S6 and phospho-Akt in both Lineage-CD45+ HSPCs and BMECs of aged mice **(Supplemental Figure 1C-D)**, confirming that aging is associated with decreased mTOR signaling within the bone marrow (BM) microenvironment.

Genetic inhibition of mTOR signaling has been demonstrated to improve lifespan and have rejuvenating effects in many model systems (Kennedy and Lamming, 2016). Indeed, it has been suggested that inhibition of mTOR signaling by Rapamycin treatment can rejuvenate aged HSPC function (Chen et al., 2009a). Contrarily, treatment of cancer patients with mTOR inhibitors is associated with infections and hematological toxicities (Rafii et al., 2015; Xu and Tian, 2014) and usage of Sirolimus in Graft-versus-host disease (GVHD) prophylaxis has been associated with hematological insufficiencies and vasculopathy (Cutler et al., 2005; Lutz and Mielke, 2016). To address these dichotomous findings, we set out to comprehensively determine the effects of mTOR inhibition on aging hematopoiesis. To this end, we treated 24 month-old mice with control feed or feed impregnated with 14 PPM Rapamycin for 2 weeks to evaluate their hematopoietic parameters **(Figure 1A)**. The 14 PPM feed has been shown to maintain plasma concentration of ∼3-5 ng/mL in C57BL6 mice that are similar to the therapeutic target levels for GVHD prophylaxis (∼3-12 ng/mL), and is considered as a ‘low dose’ regimen for mouse longevity and rejuvenation studies (Fok et al., 2014; Johnson et al., 2015; Neff et al., 2013; Zhang et al., 2014). Immunoblot analysis of WBM cells isolated from Rapamycin treated mice demonstrated a decreased expression of phospho-S6 and phospho-4EBP-1 consistent with mTOR inhibition **(Supplemental Figure 1E-F)**. The modest reduction in expression of phospho-S6 is consistent with previously reported level of inhibition in aged C57BL6 mice treated with 14 PPM diet (Zhang et al., 2014), and is likely due to the low dose of Rapamycin and the decreased basal expression of these phospho-proteins within the aged BM. Despite the modest reduction in mTOR signaling following Rapamycin, complete blood counts demonstrated that Rapamycin treated aged mice had significant decreases in white blood cell and platelet counts **(Figure 1B)**. Interestingly, the decrease in total white blood cell counts was associated with an increased frequency of myeloid cells and a decrease in T cells, likely mediated in part by the direct toxicity of Rapamycin on T cells (Molhoek et al., 2009; Tian et al., 2004) **(Figure 1C)**. Total cells per femur and the frequency of phenotypic HSPCs was not affected with Rapamycin treatment **(Figure 1D, E)**. However, functional assays for assessing progenitor potential and HSPC activity revealed significant disruption in hematopoiesis. Progenitor activity assessed by colony forming unit (CFU) assay revealed that Rapamycin treated mice displayed a significant decrease in progenitor activity **(Figure 1F)**. We next examined the long-term repopulation potential of HSPCs in a competitive BM transplantation assay, where 5 x 10^5^ whole BM (WBM) cells isolated from Rapamycin treated or control aged mice (CD45.2) were competitively transplanted along with 5 x 10^5^ CD45.1 WBM cells into lethally irradiated CD45.1 mice. BM cells from Rapamycin treated mice demonstrated a decrease in engraftment potential with a myeloid-biased output at the expense of lymphocytes, confirming a direct effect of Rapamycin at the level of HSPCs **(Figure 1G-I)**. These data suggest that systemic mTOR inhibition results in an increase in the functional defects observed in the hematopoietic system of aged mice.

**Figure 1.**
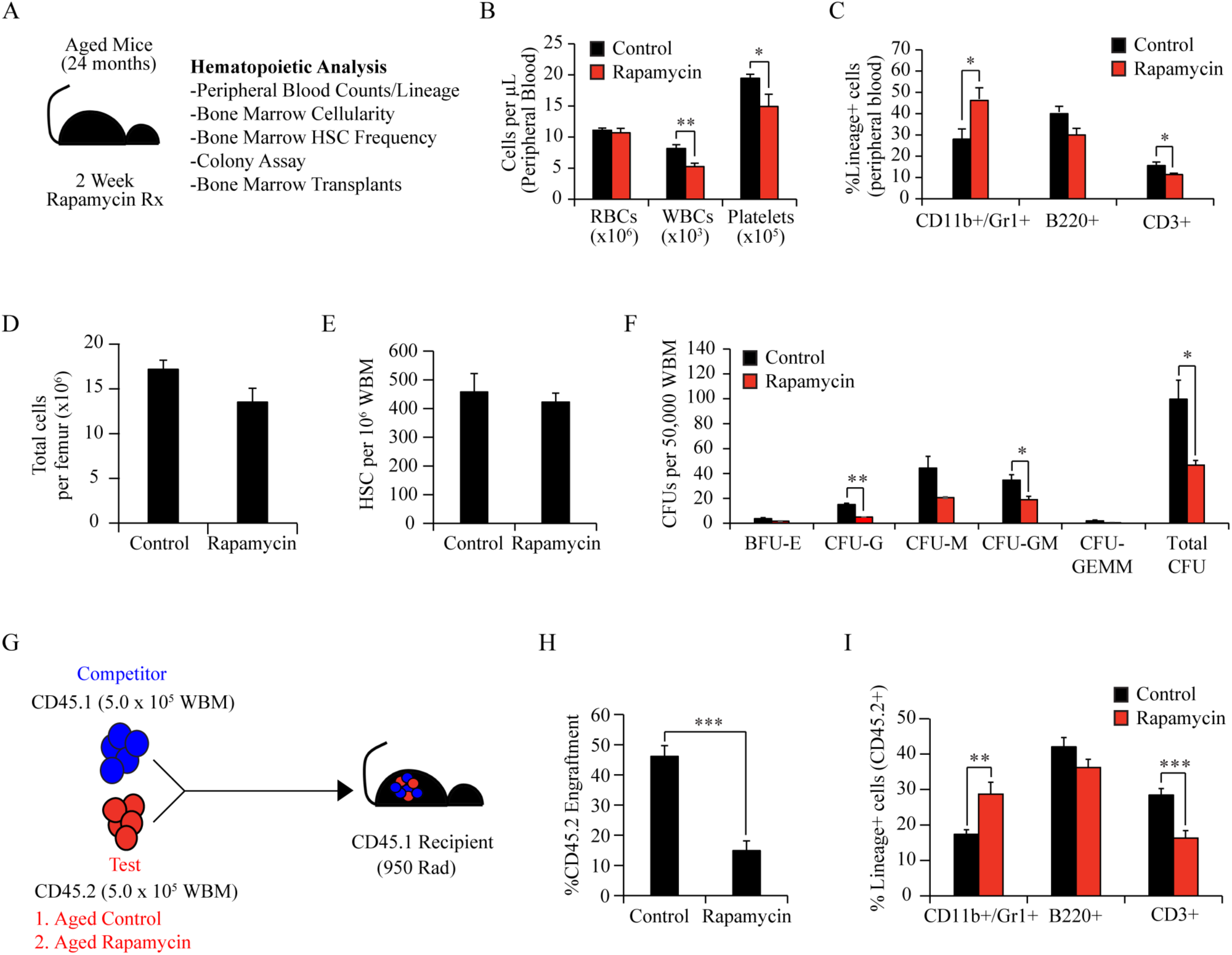
Rapamycin adversely impacts hematopoiesis and HSPC activity in aged mice. **(A)** Experimental design to assess the impact of Rapamycin on aging hematopoiesis. **(B)** Bar graphs of peripheral blood counts demonstrating a decrease in white blood cells and platelets following Rapamycin treatment (n=12 mice per cohort). **(C)** Lineage composition of peripheral blood CD45+ cells showing an increased frequency of myeloid cells in Rapamycin treated aged mice (n=8-10 mice per cohort). **(D)** Total hematopoietic cells (n=12 mice per cohort) and **(E)** the frequency of immunophenotypically-defined HSCs per femur (n=8 mice per cohort). **(F)** Methylcellulose-based colony assay by quantifying colony-forming units (CFUs) revealed a decrease in hematopoietic progenitor activity (n=3 mice per cohort). Data in **B-F** represent combined analysis of 2 independent experiments. **(G)** Schematic of competitive WBM transplantation assay to assess HSPC activity within WBM cells following Rapamycin treatment. Peripheral blood analysis after 16 weeks post-transplantation revealed a **(H)** decrease in long-term engraftment potential and **(I)** myeloid-biased lineage output at the expense of lymphopoiesis (n=10 recipients per cohort), confirming that Rapamycin adversely impacts HSPC activity in aged mice. Data represents average engraftment following transplantation of cells derived from n=3 independent donor mice per cohort. Error bars represent sample mean ± SEM. Statistical significance determined using Student’s t-test. * P<0.05; ** P<0.01; *** P<0.001.

Aberrant mTOR activation has been observed in a wide spectrum of hematological cancers as well as solid tumors and mTOR inhibitors are currently being evaluated as a stand-alone or as an adjuvant to standard chemotherapy regimens (Calimeri and Ferreri, 2017; Huang et al., 2015). The immunosuppressive properties of mTOR inhibitors are widely utilized to prevent GVHD following allogenic hematopoietic stem cell transplantation (HSCT) (Armand et al., 2016; Cutler and Antin, 2010). Chemo-radiation based conditioning regimens for HSCT as well standard induction regimens for hematologic malignancies are often associated with cytopenias resulting from the myelosuppressive injury of the endogenous HSPCs. Prior studies have shown that HSPCs undergo mTOR activation during myelosuppressive recovery (Baumgartner et al., 2018), and that mTORC1 inhibition delays hematopoietic recovery (Yan et al., 2016). To determine the impact of mTOR inhibition on regeneration of aged HSPCs, we subjected aged mice to sublethal irradiation (650 Rads) or chemotherapy (AraC 100mg/kg 5 days and Doxo 3mg/kg 3 days) and maintained the mice on either control diet or Rapamycin impregnated feed during a 28-day recovery phase **(Figure 2A, I)**. Analysis of peripheral blood revealed that Rapamycin treated mice displayed significant delay in white blood cell and platelet recovery in both injury models and a delay in neutrophil recovery following chemotherapy, indicating that mTOR signaling supports the recovery of the aged hematopoietic system following both myelosuppressive injuries **(Figure 2B, J)**. We next examined whether Rapamycin treatment impacted the number and functional potential of aged HSPCs following the 28 day hematopoietic recovery phase. Both total femur counts and the frequency of phenotypic HSPCs were significantly reduced with Rapamycin treatment suggesting that mTOR signaling is essential for HSPC recovery and restoration of BM cellularity **(Figure 2C, D, K, L)**. Additionally, Rapamycin treatment significantly affected the progenitor activity of the hematopoietic system, with a significant decrease in progenitor function following chemotherapy and radiation **(Figure 2E, M)**. To test if the decrease in phenotypic HSPCs manifests a decrease in engraftment potential, WBM (CD45.2^+^) cells were isolated from control and Rapamycin treated groups and were transplanted with a competitive dose of young CD45.1^+^ WBM in a 5:1 ratio into lethally-irradiated young CD45.1 recipients **(Figure 2F, N)**. Peripheral blood analysis after 16 weeks following transplantation revealed a significant decrease in engraftment of transplanted cells from Rapamycin treated mice from both myelosuppression groups **(Figure 2G, O)**, with Rapamycin treated mice post-irradiation resulting in no observable engraftment **(Figure 2O)**. Although Rapamycin treatment did not result in significant changes in the lineage composition of the transplanted cells **(Figure 2H, P)**, there was a notable increase in the frequency of myeloid cells **(Figure 2H)**. These data also demonstrate that irradiation (650 Rad) results in a higher degree of myelosuppressive injury as compared to the induction chemotherapy (Ara-C+Doxo) as indicated by the delayed WBC and neutrophil recovery, decreased BM cellularity and HSC frequency and engraftment potential in aged control mice receiving radiation as compared to aged control mice subject to chemotherapy. These findings are similar to previous reports demonstrating an earlier recovery of WBC counts in young mice subject to induction chemotherapy (Ara-C+Doxo) (Wunderlich et al., 2013), as compared to myelosuppressive irradiation (650 Rad)(Poulos et al., 2016; Poulos et al., 2017). Taken together, these data suggest that inhibition of mTOR signaling in aged mice results in significant hematopoietic defects at steady state, a delay in hematopoietic recovery, and in a profound impairment of HSPC function following myelosuppressive injury.

**Figure 2.**
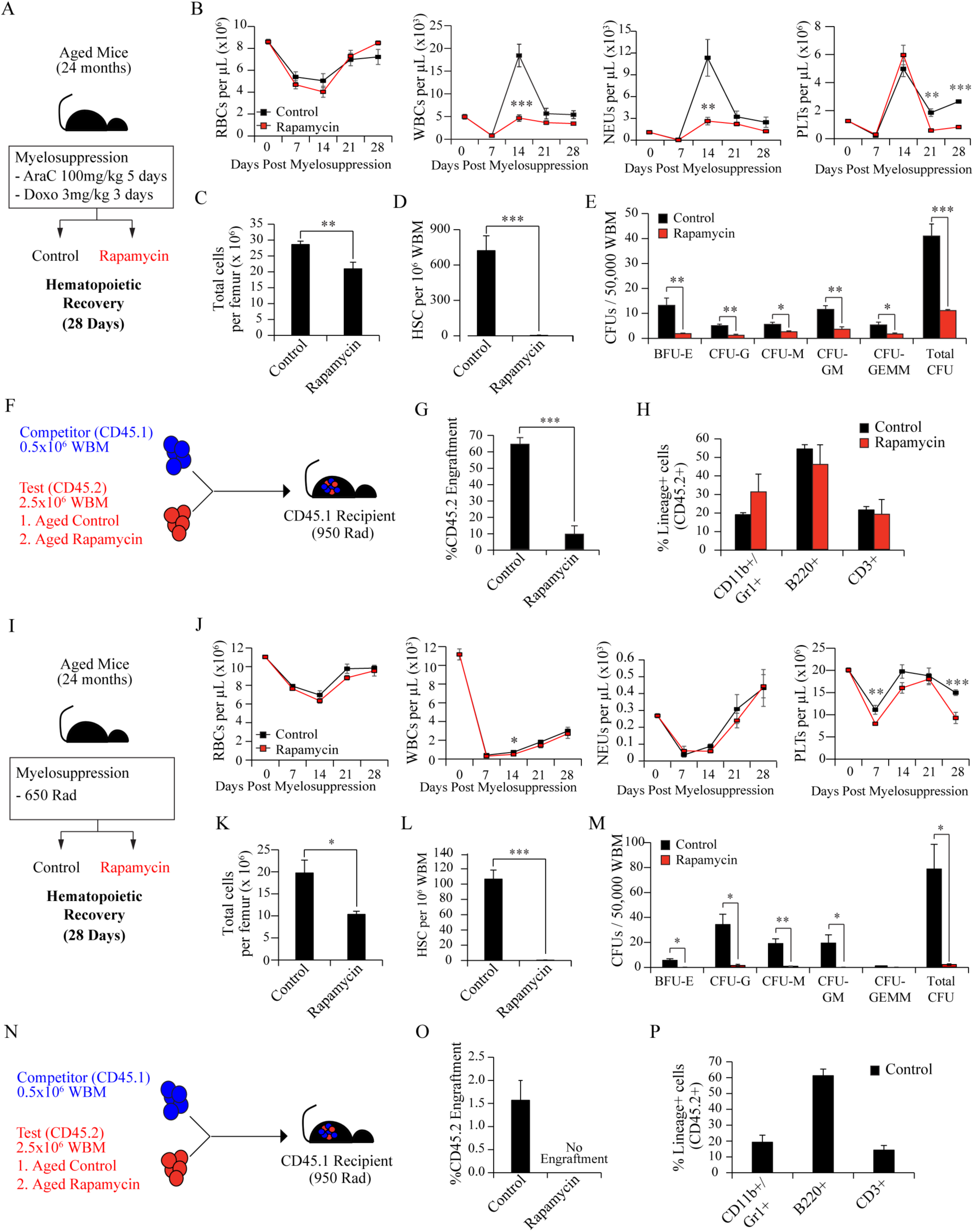
Rapamycin impairs hematopoietic recovery following myelosuppressive injury. **(A)** Experimental design to assess the impact of Rapamycin on hematopoietic recovery following chemotherapy. **(B)** Peripheral blood counts demonstrate a delay in white blood cell, neutrophil and platelet recovery following chemotherapy (n=6 mice per cohort). **(C)** Total hematopoietic cells per femur (n=5 mice per cohort) and **(D)** the frequency of HSCs per 10^6^ femur cells (n=5 mice per cohort) at day 28 following chemotherapy. **(E)** Methylcellulose-based colony assay to assess hematopoietic progenitor activity (n=3 mice per cohort). Data in **B-E** represent combined analysis of 2 independent experiments. **(F)** Schematic of competitive WBM transplantation assay to determine HSPC recovery following chemotherapy. Peripheral blood analysis after 16 weeks post-transplantation revealed a loss of (**G)** long-term engraftment potential, and **(H)** normal lineage reconstitution (n=6 recipients per cohort) in donor WBM cells derived from Rapamycin treated mice. Data represents average engraftment following transplantation of WBM cells derived from n=6 independent donor mice per cohort. **(I)** Experimental design to assess the impact of Rapamycin on hematopoietic recovery following myelosuppressive irradiation. **(J)** Peripheral blood counts demonstrate a delay in white blood cell and platelet recovery following radiation (n=11 mice per cohort). **(K)** Total hematopoietic cells per femur (n=5 mice per cohort) and **(L)** the frequency HSCs per 10^6^ femur cells (n=5 mice per cohort) at day 28 following irradiation. **(M)** Methylcellulose-based colony assay to assess hematopoietic progenitor activity (n=3 mice per cohort). Data in **J-M** represent combined analysis of 2 independent experiments. WBM transplantation assay. Peripheral blood analysis after 16 weeks post-transplantation revealed a complete loss of (**O)** long-term engraftment potential, and **(P)** multi-lineage reconstitution ability (n=4-8 recipients per cohort) in donor WBM cells derived from Rapamycin treated mice. Data represents average engraftment following transplantation of cells derived from n=4 independent donor mice per cohort. Error bars represent sample mean ± SEM. Statistical significance determined using Student’s t-test. * P<0.05; ** P<0.01; *** P<0.001.

The adverse effects of Rapamycin on hematopoiesis of aged mice likely arises due to a direct effect of mTOR inhibition within HSPCs (Guo et al., 2013) and potentially due to its additional impact on the BMEC niche function. Indeed, it has been reported that Rapamycin can lead to long-term endothelial dysfunction in patients receiving sirolimus-eluting stents (Hofma et al., 2006) and vasculopathy is a limiting toxicity in patients undergoing GVHD prophylaxis following allogenic HSCT (Cutler et al., 2005; Lutz and Mielke, 2016). However, the impact of mTOR inhibition on the ability of the endothelial niche to support HSPC activity has not been studied. Given that aged mice manifest significant hematopoietic defects following Rapamycin treatment, it is likely that enhanced hematological toxicity of aged mice to Rapamycin could be partly mediated by the adverse effects of mTOR inhibition on the vascular endothelium, in addition to the direct effect on HSPCs. To test whether endothelial mTOR is essential for maintaining niche activity, we deleted *mTOR* specifically within ECs of young adult mice (12-16 weeks) by crossing a *mTOR^fl/fl^* mouse to a tamoxifen-inducible *cre* transgenic mouse driven by the adult EC-specific VEcadherin promoter (*mTOR^(ECKO)^*) (Benedito et al., 2009). To rule out the effects of *cre*-mediated toxicity, we first analyzed *Cdh5-creERT2+* mice and observed no significant alterations in their BM cellularity, HSC frequency or peripheral blood lineage composition as compared to their *cre*-negative littermate controls, indicating that endothelial-specific expression of *cre* transgene does not affect hematopoiesis **(Supplemental Figure 1G-I)**. To determine the effect of EC-specific *mTOR* deletion on the regulation of HSPCs and their progeny, we performed hematopoietic analysis on young (12-16 weeks) *mTOR^(ECKO)^* and young (12-16 weeks) control mice. 22-24 month old wild-type mice served as aged controls. *mTOR^(ECKO)^* mice displayed a significant increase in both total BM hematopoietic cells and the frequency of phenotypic long-term repopulating HSCs (LT-HSCs), similar to aged controls **(Figure 3A-B)**. Peripheral blood analysis for lineage composition revealed a significant increase in myeloid cells in young *mTOR^(ECKO)^* mice and aged controls, with decreased levels of B cells as compared to young control mice **(Figure 3C)**. Notably, heterozygote *mTOR^fl/+^ cre-ERT2^+^* mice did not manifest the HSPC aging phenotypes observed in *mTOR^(ECKO)^* mice including increased BM cellularity, HSC frequency, and myeloid-skewed peripheral blood lineage composition demonstrating that deletion of both alleles of endothelial *mTOR* are essential to induce HSPC aging phenotypes **(Supplemental Figure 1J-L)**. Next, we determined that whole bone marrow isolated from *mTOR^(ECKO)^* and aged mice displayed a decrease in progenitor activity in methylcellulose colony forming unit (CFU) assay **(Figure 3D)**. We further analyzed HSCs from *mTOR^(ECKO)^* mice for levels of *γ*H2AX foci (Flach et al., 2014) and cell polarity status (Florian et al., 2012). HSCs from *mTOR^(ECKO)^* mice and aged controls displayed a significant increase in *γ*H2AX foci and a striking loss of *α*TUBULIN polarity **(Figure 3E-H)**, as compared to young control mice. Transcriptional analysis revealed that EC-specific *mTOR* deletion leads to changes in HSC gene expression that cluster with aged HSCs **(Figure 3I)**. To define a robust gene expression signature that characterizes an aged HSC, we compared our microarray data with prior datasets published by *Rossi et al* (Rossi et al., 2005) and *Chambers et al* (Chambers et al., 2007) and identified ten genes that show significant upregulated expression with aging **(Figure 3J)**. RT-qPCR analysis confirmed an upregulation of all of the ten ‘aging-signature’ genes in aged-HSCs **(Figure 3K)**. Strikingly, *mTOR^(ECKO)^* HSCs also displayed a similar upregulation of the ‘aging-signature’ genes, as observed in aged HSCs **(Figure 3K)**. Taken together, these observations suggest that EC-specific *mTOR* deletion is sufficient to induce premature aging of the HSC and the hematopoietic system during homeostasis.

**Figure 3.**
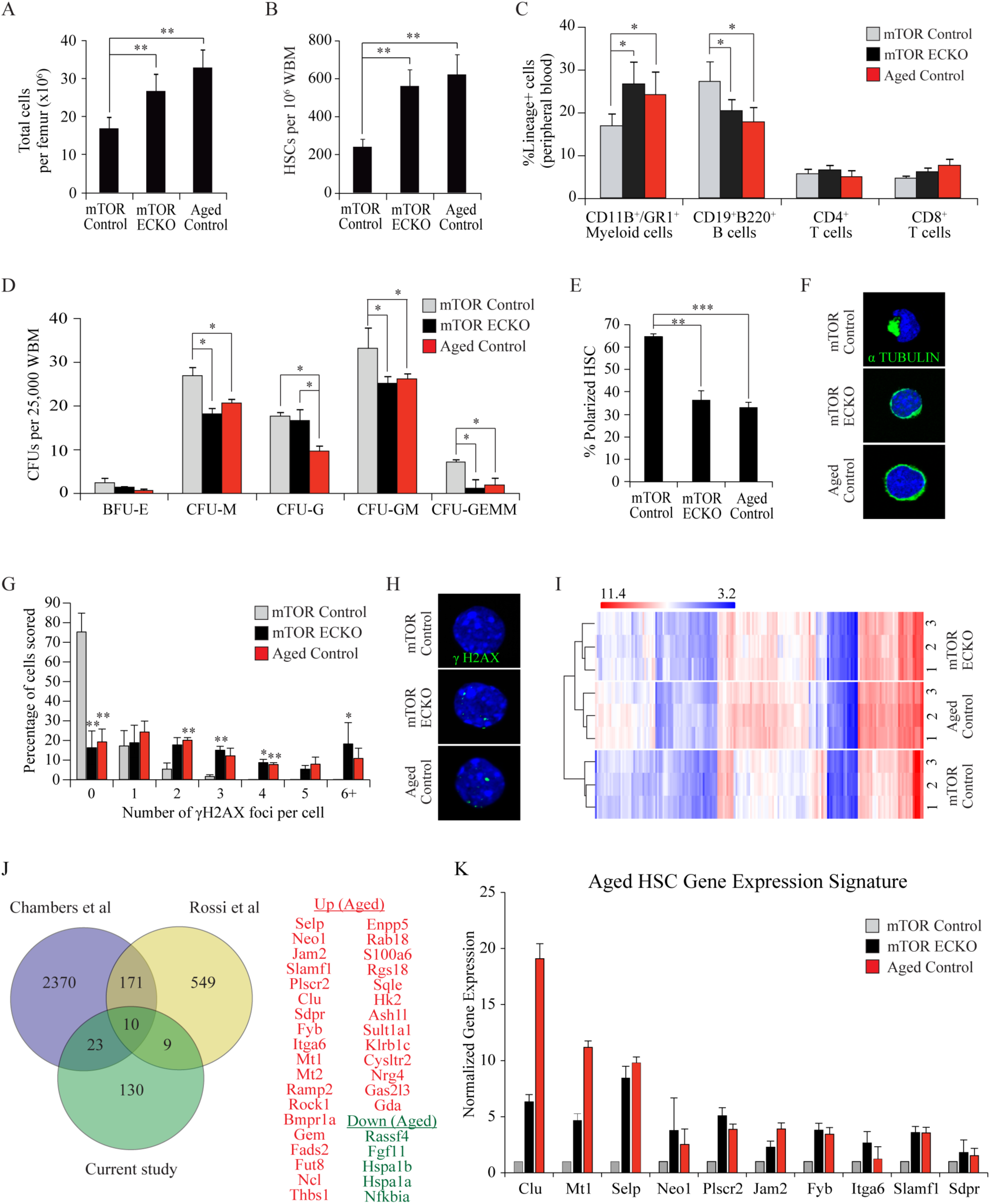
EC-specific deletion of *mTOR* in young mice leads to aging-related alterations in HSPCs. To determine the effect of endothelial-specific *mTOR* deletion on HSPCs, hematopoietic analysis **(A-C)** was performed on young *mTOR^fl/fl^(n=8)*, *mTOR^(ECKO)^ (n=10),* and aged control mice *(n=5)*. **(A, B)** Endothelial-specific deletion of *mTOR* resulted in **(A)** a significant increase in total hematopoietic cells and **(B)** the frequency of immunophenotypically-defined LT-HSCs per femur. **(C)** Lineage composition of CD45+ hematopoietic cells in the peripheral blood. **(D)** Methylcellulose-based colony assay was used to assess hematopoietic progenitor activity by quantifying colony-forming units (CFUs) (n=3 mice per cohort). **(E)** Quantification of the frequency of polarized HSCs (n=3 mice per cohort). **(F)** Representative images of *α*TUBULIN staining to evaluate HSC cellular polarity. **(G)** Quantification of number of *γ*H2AX foci per HSC (n=3 mice per cohort). **(H)** Representative images of *γ*H2AX foci in HSC. Data in **A-H** represent combined analysis of 2 independent experiments. **(I)** Transcriptional profiling of HSCs by microarray in the indicated genotypes (n=3 mice per cohort). Note: The phenotypes observed in *mTOR^(ECKO)^* mice closely mimic the phenotypes observed in aged control mice. **(J)** Venn Diagram comparing genes showing significant changes between young and aged hematopoietic stem cells in the current study with two previously published datasets. Genes listed demonstrate concordant changes in expression between the current study and published datasets (red - upregulated in aged HSCs; green - downregulated in aged HSCs). Genes in bold text were validated by RT-qPCR and represent an aged HSC expression signature. **(K)** RT-qPCR confirmation of microarray-identified aged HSC gene expression signature in *mTOR^(ECKO)^* and aged mice (n=3 mice per cohort). Note: HSCs from *mTOR^(ECKO)^* mice share an aged HSC gene expression signature. Error bars represent mean ± SEM. Statistical significance determined using One-way ANOVA with Tukey’s Correction for multiple comparisons. *P<0.05, **P<0.01, ***P<0.001.

To verify whether these age-related alterations in *mTOR^(ECKO)^* mice are due to direct effects on the HSC, we examined the long-term repopulation capacity of HSCs in a BM transplantation assay. CD45.2^+^ HSCs from young *mTOR^(ECKO)^*, young control and aged mice were competitively transplanted into lethally-irradiated CD45.1 mice **(Figure 4A)**. HSCs from young *mTOR^(ECKO)^* mice displayed diminished engraftment and a significant myeloid bias at the expense of T-lymphopoiesis **(Figure 4B)**, as compared to HSCs from young control mice. Notably, the functional attributes of *mTOR^(ECKO)^* HSCs are similar to HSCs from aged mice. To determine whether HSCs from *mTOR^(ECKO)^* mice display self-renewal defects, we performed secondary transplantation assays **(Figure 4C)**. Additionally, we examined whether a BM endothelial niche devoid of mTOR signaling is sufficient to drive young control HSPCs from primary transplants to functionally behave like aged HSPCs **(Figure 4C)**. To this end, we isolated and transplanted WBM from primary recipients (that received 100 HSCs from either young control or *mTOR^(ECKO)^* mice) into lethally-irradiated young control or young *mTOR^(ECKO)^* secondary recipients **(Figure 4C)**. WBM from aged-HSC primary recipients was transplanted into young wild type secondary recipients as an aged secondary transplant control **(Figure 4D-G; red dashed line represents average values)**. As expected, WBM isolated from young control primary recipients, when transplanted into secondary control recipients **(Figure 4D-G; CNTL:CNTL),** maintained consistent overall long-term engraftment and balanced myelo-lymphoid output. As anticipated, WBM from aged-HSC primary recipients when transplanted into secondary recipients displayed further decrease in overall engraftment coupled with a myeloid biased output **(Figure 4D-G; red dashed line)**. However, WBM isolated from young control primary recipients, when transplanted into secondary *mTOR^(ECKO)^* recipients **(Figure 4D-G; CNTL:ECKO)** resulted in a significant decrease in CD45.2 engraftment, with the development of a myeloid bias and a decrease in T cell production, confirming that EC-specific deletion of *mTOR* is sufficient to impose aging attributes on HSPCs derived from young control mice. Conversely, transplantation of primary *mTOR^(ECKO)^* WBM into control secondary recipients **(Figure 4D; ECKO:CNTL)**, led to a modest increase in CD45.2 engraftment potential as compared to the primary *mTOR^(ECKO)^* WBM that was transplanted into *mTOR^(ECKO)^* recipients **(Figure 4D; ECKO:ECKO)**. Although a myeloid bias and a decreased T cell output still persisted in the ECKO:CNTL cohort **(Figure 4E, G)**, transplanting primary *mTOR^(ECKO)^* WBM into control recipients (ECKO:CNTL) resulted in a significant reduction in the percentage of myeloid cells and in an increase in T cells as compared to the ECKO:ECKO cohort **(Figure 4E, G),** suggesting that exposure to a control microenvironment partially restores hematopoietic function. When primary *mTOR^(ECKO)^* WBM was transplanted into secondary *mTOR^(ECKO)^* recipients (Fig. 4B; ECKO:ECKO), the engraftment was similar to the aged secondary control cohort **(Figure 4D; red dashed line)**, with a persisting myeloid bias and decreased T cell output. These data demonstrate that primary control HSPCs can be instructed to assume aging phenotypes following transplantation when introduced into a *mTOR^(ECKO)^* microenvironment and that partial reversal of the aging phenotypes observed in *mTOR^(ECKO)^* HSPCs transplanted from primary recipients can be achieved when introduced into a wild-type BM microenvironment.

**Figure 4.**
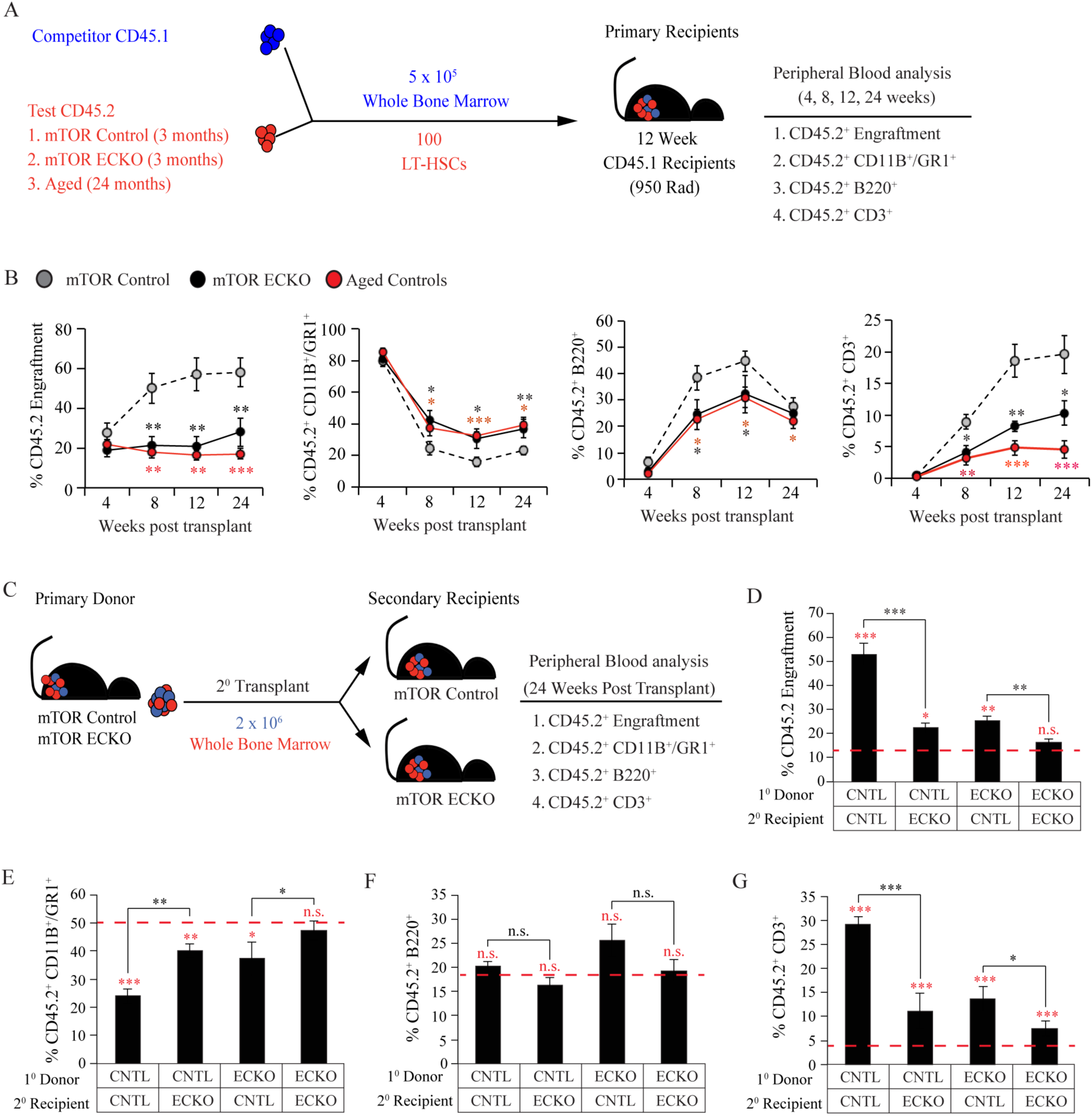
HSCs from young *mTOR^(ECKO)^* mice display functional defects resembling aged HSCs. **(A)** Schematic of primary transplants. **(B)** Long-term, multi-lineage repopulation capacity of HSCs (n=10 recipients per cohort). CD45.2^+^ HSCs transplanted from young *mTOR^(ECKO)^* mice displayed diminished engraftment potential and a significant myeloid bias at the expense of lymphopoiesis. Data represents average engraftment following transplantation of cells derived from n=3 independent donor mice per cohort. **(C)** Schematic of secondary transplants. Whole bone marrow isolated from primary recipients were transplanted into either young *mTOR^fl/fl^* control or young *mTOR^(ECKO)^* mice. Primary recipients that received aged HSCs were transplanted into young wild type secondary recipient mice and served as aged controls. **(D-G)** Quantification of multi-lineage engraftment in secondary transplant recipients. **(D)** CD45.2^+^ peripheral blood engraftment and **(E)** myeloid, **(F)** B cell, and **(G)** T cell lineage-committed contribution after 16 weeks post-transplantation (n=5-7 recipients per cohort). Data represents average engraftment following transplantation of cells derived from n=5 independent donor mice per cohort. Error bars represent mean ± SEM. Statistical significance determined using Student’s t-test. *P<0.05, **P<0.01, ***P<0.001, n.s. denotes not statistically significant.

We next sought to evaluate the roles of mTORC1 and mTORC2 complexes downstream of endothelial mTOR in HSPC regulation. To this end, we generated bone marrow-derived endothelial cell lines deficient for Raptor (mTORC1) or Rictor (mTORC2), and utilized our established co-culture platform (Poulos et al., 2015) to test their impact on HSPC activity **(Supplemental Figure 2)**. The functional knockdown of either complex was confirmed by immunoblot analysis for downstream targets of Raptor and Rictor which demonstrated a decrease in phospho-4EBP-1 upon knockdown of Raptor and a decrease in phospho-Akt upon Rictor knockdown **(Supplemental Figure 2A-G)**. Analysis after 9 days of co-culture revealed no significant differences in total hematopoietic expansion after knockdown of either complex **(Supplemental Figure 2H-I)**. However, colony assays revealed a significant reduction in CFU-forming ability of HSPCs co-cultured on Raptor as well as Rictor knockdown endothelial cells **(Supplemental Figure 2J)**. Notably, the total number of CFU-GEMMs were significantly decreased after knockdown of either Raptor or Rictor, suggesting a non-redundant role of endothelial mTORC1 and mTORC2 in maintaining HSPC function. Whether each of these complexes regulate distinct attributes of HSPC aging phenotypes *in vivo* remains to be addressed.

Given the adverse impact of systemic mTOR inhibition on hematopoietic recovery **(Figure 2)**, we next sought to determine whether EC-specific deletion of *mTOR* influences post-myelosuppressive hematopoietic reconstitution and HSPC recovery. We subjected *mTOR^(ECKO)^* and littermate controls to myelosuppressive injury (650 Rad) and assessed their hematopoietic recovery **(Supplemental Figure 3A)**. Analysis of peripheral blood revealed that *mTOR^(ECKO)^* mice displayed a significant delay in hematopoietic recovery indicating that endothelial mTOR signaling is essential for efficient recovery following myelosuppressive injury **(Supplemental Figure 3B)**. After allowing 28 days for recovery, we assessed total marrow counts per femur and the frequency of phenotypic HSCs and found no significant differences between control and *mTOR^(ECKO)^* mice **(Supplemental Figure 3C)**. However, progenitor activity, particularly the CFU-GEMM and Total CFU counts, was significantly impaired in BM cells derived from *mTOR^(ECKO)^* mice **(Supplemental Figure 3E)**. We next examined the functional output of the HSPCs of *mTOR^(ECKO)^* mice at day 28 post myelosuppression by transplanting CD45.2^+^ donor cells from control and *mTOR^(ECKO)^* mice along with a competitive dose of young CD45.1^+^ WBM in a 5:1 ratio into lethally-irradiated young CD45.1 recipients **(Supplemental Figure 3F)**. Long-term, multi-lineage engraftment was analyzed four months post-transplantation and we found that EC-specific deletion of mTOR resulted in profound defects in engraftment potential following myelosuppressive injury along with an impaired frequency of CD3+ T cells **(Supplemental Figure 3G-H)**. Collectively, this data demonstrates deletion of mTOR signaling within the endothelial niche results in hematopoietic defects that resemble the defects observed during Rapamycin treatment. Additionally, these observations suggest that EC-specific *mTOR* deletion in young mice is sufficient to induce transcriptional, phenotypic, and functional aging of the HSCs at steady state and during regeneration.

To obtain insights into the mechanisms by which endothelial mTOR influences HSPC aging phenotypes, we studied the impact of mTOR deletion on phenotypic and functional characteristics of the bone marrow endothelium **(Figure 5)**. Deletion of mTOR within endothelium of young mice did not manifest any gross changes in BM vascular morphology **(Figure 5A)**, total number of endothelial cells within the bone marrow **(Figure 5B)** or their apoptosis **(Figure 5C)**. mTOR deletion, however, was associated with a modest increase in percentage of endothelial cells in G0 phase of cell-cycle **(Figure 5D)**, consistent with the positive role of mTOR in cell-cycle progression. Notably, *mTOR^(ECKO)^* mice do not manifest any gross defects in their survival which is in stark contrast to the deletion of mTOR within hematopoietic cells of adult mice wherein the mice die within ∼2 weeks due to hematopoietic failure (Guo et al., 2013). These findings illuminate the differential tissue-specific roles of mTOR in adult mice; while HSPCs are constantly cycling in order to replenish mature peripheral blood cell output and hence dependent on active mTOR signaling for cell growth and division, endothelial cells are predominantly quiescent in the adult bone marrow (∼ 1% of cells are in SG2M phase). These findings also likely explain the relatively higher degree of toxicity observed with Rapamycin during myelosuppressive recovery as compared to steady state situations wherein active mTOR signaling is likely to be essential for efficient recovery of both HSPCs and endothelial cells. Collectively, these findings indicate that the HSPC aging phenotypes observed in young *mTOR^(ECKO)^* mice do not arise from gross vascular disruption but rather likely due to changes in the instructive signals originating from the endothelial niche.

**Figure 5.**
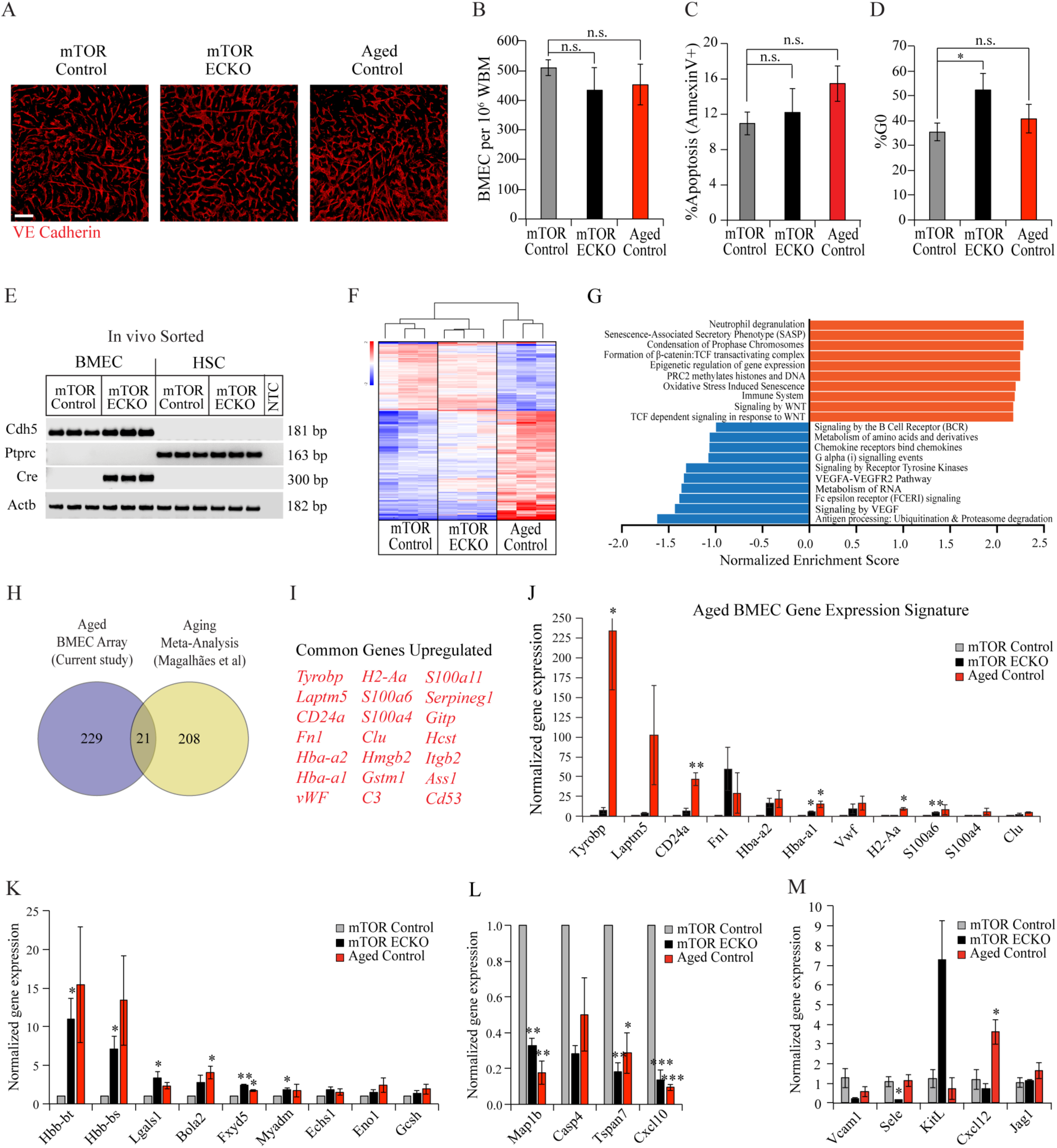
Loss of endothelial mTOR recapitulates a subset of transcriptional changes observed during physiological ageing. **(A)** Representative immunofluorescence images of femurs intravitally-labeled with a vascular-specific VECAD antibody (red) demonstrating normal vascular morphology in *mTOR^(ECKO)^* mice (n=4 mice/cohort). Scale bar represents 100 µm **(B)** Frequency of phenotypic BMECs per 10^6^ femur cells assessed by flow cytometry (n=7 mice/cohort) **(C)** Frequency of apoptotic BMECs (Annexin V+) assessed by flow cytometry (n=7 mice/cohort). **(D)** Cell-cycle analysis of BMECs (Ki-67/Hoechst staining) assessed by flow cytometry (n=7 mice/cohort). **(E)** Agarose gel electrophoresis image of RT-PCR amplicons for the indicated genes using RNA isolated from FACS sorted BMECs and HSCs in the indicated genotypes (n=3 mice per cohort). NTC denotes ‘No Template Control’. Data in **A-E** represent combined analysis of 2 independent experiments. **(F)** Transcriptional profiling of BM ECs by microarray in the indicated genotypes (n=3 mice per cohort). **(G)** Reactome Pathway analysis of genes differentially expressed in aged BMECs as compared to young BMECs. **(H)** Venn Diagram comparing genes showing significant changes between aged BM ECs in the current study with a previously published meta-analysis. **(I)** Genes listed demonstrate concordant changes in expression between the current study and published dataset. **(J)** RT-qPCR evaluation of aging-signature genes in BMECs derived from *mTOR^(ECKO)^* and aged mice. (n=3 mice per cohort). Note: BMECs from *mTOR^(ECKO)^* mice demonstrate gene expression changes similar to aged BMECs. **(K-L)** RT-qPCR evaluation of genes identified to exhibit congruent expression in BMECs derived from *mTOR^(ECKO)^* and aged mice by microarray (n=3 mice per cohort). **(M)** RT-qPCR evaluation of known angiocrine factors in BMECs derived from *mTOR^(ECKO)^* and aged mice (n=3 mice per cohort). Error bars represent sample mean ± SEM. Statistical significance determined using Student’s t-test. * P<0.05; ** P<0.01; *** P<0.001, n.s. denotes not statistically significant.

To address this possibility, we performed transcriptomic analysis on bone marrow endothelial cells derived from young control mice, young *mTOR^(ECKO)^* and aged control mice. The purity of FACS sorted cells was confirmed by RT-PCR analysis **(Figure 5E)** for expression of *Cdh5* (endothelial identity) and absence of expression of *Ptprc* (hematopoietic identity). We also confirmed the fidelity of the cre-ERT2 mouse model by performing RT-PCR analysis for the *creERT2* transgene which demonstrated robust expression within endothelial cells and was not detected in FACS-sorted HSCs **(Figure 5E)**. Unsupervised hierarchical clustering of the microarray data revealed that aged endothelial cells demonstrate a unique transcriptional profile as compared to young control mice and young *mTOR^(ECKO)^* mice **(Figure 5F)**. This indicated that, in addition to decreased mTOR signaling, aging is likely associated with changes in multiple signaling pathways and molecular processes within endothelial cells. To gain insights into these processes, we performed a Pathway analysis of differentially expressed genes in the aged BM endothelium, which revealed an over-representation of senescence associated secretory pathway (SASP) and oxidative stress along with a decrease in ubiquitin-proteasome processing and amino acid metabolism pathways **(Figure 5G)**. Interestingly, there was an increase in WNT signaling pathway which has recently emerged as a regulator of the aging process (Gruber et al., 2016), and a decrease in VEGF and tyrosine kinase signaling pathways that have been well established to play key roles in regulating endothelial signaling, and likely explain the decreased mTOR signaling in aged BM ECs. To define an aging-associated gene expression signature of BM endothelium, we compared the genes that were differentially expressed in aged endothelial cells in our microarray data with a previously published meta-analysis of aging-associated genes (de Magalhaes et al., 2009) **(Figure 5H)** and identified 21 genes that were commonly upregulated **(Figure 5I)**. Notably, *Hba* has recently been demonstrated to play a key role in regulation of endothelial nitric oxide signaling (Straub et al., 2012) which could potentially impact HSPC function through regulation of shear stress (Pardanaud and Eichmann, 2009). We next compared the expression of these endothelial aging-signature genes within BM ECs derived from young control mice, young *mTOR^(ECKO)^* and aged control mice by RT-qPCR analysis **(Figure 5J)**. Interestingly, most of these aging-associated signature genes including *Fn1*, *vWF*, *Hba-a1*, *Hba-a2*, *S100a6* and *Clu* demonstrated increased expression in endothelial cells derived from *mTOR^(ECKO)^* mice, similar to aged mice **(Figure 5J)**. This led to the possibility that aberrant expression of genes commonly altered in endothelial cells of *mTOR^(ECKO)^* mice and aged mice, as compared to young endothelial cells, could likely reveal candidates that mediate aging-associated HSPC defects. Further analysis of our transcriptome data set revealed 13 such genes **(Figure 5K-L)**, including genes implicated in regulation of cell-matrix interactions (*Lgals1*, *Fxyd5*), metabolism (*Bola2, Echs, Eno1*, *Gcsh*) and myeloid differentiation of HSPCs (*Myadm*). *Myadm* has recently been shown to regulate endothelial inflammation (Aranda et al., 2013) and increased expression of *Myadm* was observed in mesenchymal stromal cells (MSCs) derived from patients of myeloproliferative neoplasms (Ramos et al., 2017). Since inflammation-associated aging or ‘inflammaging’ has recently emerged as a critical process driving HSPC aging phenotypes, it is likely that genes regulating endothelial inflammation might impact HSPC aging (Kovtonyuk et al., 2016). We also compared the expression of angiocrine factors that are known to regulate HSC function and did not observe any congruent changes in their expression in *mTOR^(ECKO)^* mice and aged mice **(Figure 5M)**. Collectively, the genes identified by microarray analysis likely represent novel regulators of endothelial niche-HSC interactions and need to be investigated further.

Physiological aging of the HSPC microenvironment results in a decline in the functional potential of HSPCs to reconstitute hematopoiesis, thereby leading to an imbalance of the myeloid/lymphoid ratio, with the skewing towards the myeloid lineage (Ergen et al., 2012; Guidi et al., 2017; Vas et al., 2012). Tipping the balance towards myeloid output can lead to a multitude of age-associated defects, including a significant decline in the body’s ability to mount an adaptive immune response coupled with a predisposition towards myeloid neoplasms. Recent studies have indicated the possibility that systemic factors can contribute towards HSPC aging (Ju et al., 2007; Song et al., 2012) and that exposure of aged HSPCs to a young microenvironment can partially rejuvenate their defects (Yamamoto et al., 2018). It has also been demonstrated that aged BMECs have reduced function in regards to regulating hematopoietic niche cells and that it is possible to rejuvenate their functional readout; however, these studies were unable to restore the full functional capacity of the HSPC (Kusumbe et al., 2016). The observation of decreased mTOR signaling within the endothelium of aged mice raises the possibility that activation of mTOR signaling specifically within endothelial cells could alleviate the aging associated HSPC defects. Future advances in the ability to selectively target the vascular endothelium with pharmacological agents would enable targeted delivery of molecules that activate endothelial mTOR signaling during the aging process, likely leading to improved hematopoietic outcomes.

## Concluding Remarks

Here, the data presented support the hypothesis that aged BMECs undergo alterations affecting their instructive capacity to properly balance the rate of HSPC maintenance, self-renewal and differentiation. In particular, changes in BMEC signaling pathways likely contribute to the aging phenotypes seen in the hematopoietic system. This study demonstrates that the AKT/mTOR axis within ECs is critical in preserving HSPC function and that disruption of mTOR signaling specifically in ECs drives pathological phenotypes reminiscent of the hematopoietic defects observed during physiological aging. Furthermore, these data indicate that dissecting the role ECs play in modulating HSPC activity can lead to a better understanding of endothelial contributions in the initiation and progression of many of the common age-related hematopoietic disorders.

Additionally, our data indicate that pharmacological inhibition of mTOR signaling using Rapamycin, a widely regarded rejuvenating agent and clinically utilized immunosuppressant, has deleterious effects on the aging hematopoietic system, potentially due its adverse impact on the endothelial niche. Aberrant mTOR activation has been observed in a multitude of cancers and mTOR inhibitors are currently being considered as adjuvant agents for a wide array of hematological malignancies based on promising pre-clinical evidence (Calimeri and Ferreri, 2017). However, a recent large-scale clinical trial evaluating the efficacy of mTOR inhibitor Everolimus as an adjuvant to post-induction chemotherapy in improving relapse-free survival in AML patients was terminated on account of excess mortality in the Everolimus cohort (Burnett et al., 2018). Our results indicate that mTOR inhibition has a deleterious impact on hematopoietic recovery following radiation and chemotherapy-induced myelosuppressive injury in aged mice. Collectively, these observations indicate that hematologic toxicities of mTOR inhibitors must be carefully considered during evaluation of the potential therapeutic benefit of mTOR inhibitors in rejuvenation strategies targeted towards the elderly patient population.

## Materials and Methods

### Animal use

All murine experiments were conducted in accordance with the Association for Assessment and Accreditation of Laboratory Animal Care, Intl. (AAALAC) and National Institutes of Health (NIH) Office of Laboratory Animal Welfare (OLAW) guidelines and under the approval of the Center for Discovery and Innovation and the Institutional Animal Care and Use Committee (IACUC). Young and aged C57BL/6 (CD45.2) mice were purchased from the National Institute on Aging and The Jackson Laboratory. Congenic C57BL/6J *mTOR^fl/fl^* mice (B6.129S4-*Mtor^tm1.2Koz^*/J) (Jax Stock No: 011009) were purchased from The Jackson Laboratory and used as described. These mice have been backcrossed to C57BL/6 mice for 10 generations by the donating investigator prior to submission at The Jackson Laboratory. *mTOR^fl/fl^* mice were maintained by crossing with congenic C57BL/6 mice obtained from The Jackson Laboratory (Jax Stock No. 000664). C57BL/6 Tg(*Cdh5-cre/ERT2*)1Rha mice were obtained from Ralf Adams and the model was generated in a C57BL/6 background (Benedito et al., 2009). Founder lines were backcrossed to C57BL/6 by the donating investigator for at least five generations. C57BL/6 Tg(Cdh5-cre/ERT2)1Rha mice were maintained by crossing with C57BL/6 mice obtained from The Jackson Laboratory (Jax Stock No. 000664). To generate endothelial-specific *mTOR* knockout mice (*mTOR^(ECKO)^*), *mTOR^fl/fl^* mice were crossed with *Cdh5-creERT2* mice. *mTOR^fl/fl^*; *Cdh5-creERT2* mice (8-12 weeks) were administered 200 mg/kg tamoxifen (Sigma-Aldrich) via intraperitoneal injection at a concentration of 30 mg/mL in sunflower oil (Sigma-Aldrich) on three consecutive days, followed by three days off, and three additional days of injection. *mTOR^fl/fl^* Cre-negative littermate mice (8-12 weeks) were utilized as ‘young mTOR controls’. Aged C57BL/6 mice (18-20 months) were purchased from the National Institute on Aging and The Jackson Laboratory and were allowed to acclimatize in our vivarium for at least 1 month prior to experiments, and were utilized as ‘aged controls’. Young and aged control mice received parallel tamoxifen administration along with *mTOR^(ECKO)^*) mice. Experiments were conducted at least one month post-tamoxifen administration. B6.SJL-*Ptprc^a^Pepc^b^*/BoyJ (CD45.1) mice were purchased from The Jackson Laboratory and used as described. For transplantation studies, recipient mice were subjected to 950 Rad of total body irradiation (*^137^Cs*) 24 hours prior to transplantation. Transplant recipients were maintained PicoLab Mouse 20 antibiotic feed (0.025% Trimethoprim and 0.124% Sulfamethoxazole; LabDiet) for 28 days following irradiation. Microencapsulated Rapamycin feed was prepared according to a previously described protocol (Harrison et al., 2009).

### Buffers and Media

1. Magnetic activated cell sorting (MACS) buffer: PBS without Ca^++^/Mg^++^ (pH 7.2) (Corning 21-040-CV) containing 0.5% bovine serum albumin (BSA; Fisher Scientific BP1605) and 2 mM EDTA (Corning 46-034-CI).
2. Digestion buffer: 1x Hanks Balanced Salt Solution (Life Technologies 14065) containing 20 mM HEPES (Corning 25-060-CI), 2.5 mg/mL Collagenase A (Roche 11088793001), and 1 unit/mL Dispase II (Roche 04942078001).
3. Endothelial growth medium: Low-glucose DMEM (ThermoFisher Scientific 11885-084) and Ham’s F-12 (Corning 10-080) (1:1 ratio), supplemented with 20% heat-inactivated FBS (Denville Scientific FB5002-H), 1% antibiotic-antimycotic (Corning 30-004-CI), 1% non-essential amino acids (Corning 25-025-CI), 10 mM HEPES (Corning 25-060-CI), 100 µg/mL heparin (Sigma-Aldrich H3149), and 50 µg/mL endothelial cell growth supplement (Alfa Aesar BT-203)].

### Peripheral blood analysis

Peripheral blood (PB) was collected using 75 mm heparinized glass capillary tubes (Kimble-Chase) via retro-orbital sinus bleeds into PBS (pH 7.2) containing 10 mM EDTA. WBC, Neutrophil, RBC, and platelet counts were analyzed using an Advia120 (Bayer Healthcare). To quantify steady state lineage^+^ hematopoietic cells and mutli-lineage HSC engraftment, PB was depleted of red blood cells (RBC Lysis Buffer; Biolegend) according to the manufacturer’s recommendations, stained with indicated fluorophore-conjugated antibodies, and analyzed using flow cytometry.

### WBM analysis

To quantify total hematopoietic cells, femurs were gently crushed with a mortar and pestle and enzymatically disassociated for 15 min at 37°C in Digestion buffer following which cell suspensions were filtered (40 µm; Corning 352340) and washed in MACS buffer. Viable cell numbers were quantified using a hemocytometer with Trypan Blue (Life Technologies) exclusion. To quantify hematopoietic stem and progenitor cells (HSPCs) in the BM, femurs were flushed using a 26.5-gauge needle with MACS buffer. To quantify bone marrow endothelial cells (BMECs, defined as CD45-CD31+ VEcadherin+ cells), mice were injected retro-orbitally with 25 µg of a fluorophore-conjugated antibody raised again VE-cadherin (BV13; Biolegend) 15 minutes prior to sacrifice. Femurs were gently crushed with a mortar and pestle and enzymatically disassociated for 15 min at 37°C in Digestion buffer following which cell suspensions were filtered and washed in MACS buffer. Cells were surface stained using fluorochrome-conjugated antibodies as per manufacturer recommendations. Cell populations were analyzed using flow cytometry.

### Flow cytometry and cell sorting

Prior to surface staining, F_c_ receptors were blocked using a CD16/32 antibody (93; Biolegend) in MACS buffer for 10 minutes at 4°C. F_c_-blocked samples were stained with fluorophore-conjugated antibodies at manufacturer-recommended concentrations in MACS buffer for 30 minutes at 4°C. Stained cells were washed in MACS buffer and fixed in 1% paraformaldehyde (PFA) in PBS (pH 7.2) with 2 mM EDTA for flow cytometric analysis or resuspended in PBS (pH 7.2) with 2 mM EDTA and 1 µg/mL 4-6, Diamidino-2-Phenylindole (DAPI) (Biolegend) for live/dead exclusion and cell sorting. Samples were analyzed using a LSR II SORP (BD Biosciences) and sorted using an ARIA II SORP (BD Biosciences). Data was collected and analyzed using FACS DIVA 8.0.1 software (BD Biosciences).

### Phosphoflow cytometry

To assess mTOR phosphorylation, young (12-16 weeks) and aged (24 months) C57BL6 mice were injected retro-orbitally with 25 µg of a fluorophore-conjugated antibody raised again VE-cadherin (BV13; Biolegend) 15 minutes prior to sacrifice. Long bones were isolated, crushed, and enzymatically disassociated in Digestion buffer for 15 minutes at 37°C. Resulting cell suspensions were filtered (40 µm), washed using MACS buffer. Single-cell WBM suspensions were depleted of lineage-committed hematopoietic cells using a Lineage Cell Depletion Kit (Miltenyi) according to the manufacturer’s suggestions. Resulting lineage^-^ cells were stained with fluorophore-conjugated antibodies raised against CD31 (390; Biolegend) and CD45 (30-F11; Biolegend). Stained cells were washed using MACS buffer, fixed, permeabilized using Phosphoflow Fix Buffer 1 and Perm Buffer 3 (BD Biosciences) and stained with antibodies raised against phosphorylated mTOR (Ser2448) (BD Biosciences 563489), phosphorylated AKT (S473) (BD Biosciences 560404) and phosphorylated S6 (S235/236) (BD Biosciences 560434) for 30 minutes at room temperature according to the manufacturer’s recommendations. Cells were washed in MACS buffer. Appropriate concentration matched isotype controls were utilized for gating and analysis by Flow cytometry.

### Cell cycle and apoptosis

Mice were injected retro-orbitally with 25 µg of a fluorophore-conjugated antibody raised again VE-cadherin (BV13; Biolegend) 15 minutes prior to sacrifice. Long bones were isolated, crushed, and enzymatically disassociated in Digestion buffer for 15 minutes at 37°C. Resulting cell suspensions were filtered (40 µm), washed using MACS buffer. Single-cell WBM suspensions were surface-stained with fluorophore-conjugated antibodies raised against CD31 (390; Biolegend) and CD45 (30-F11; Biolegend). Stained cells were washed in MACS buffer. For cell cycle analysis of BMECs, surface stained cells were fixed and permeabilized using the BD Cytofix/Cytoperm Kit (BD Biosciences) as per manufacturer’s suggestions following which cells were stained with an antibody raised against Ki67 (B56, BD 558616) and counterstained with Hoechst 33342 (BD Biosciences), according to the manufacturer’s recommendations. Cell were analyzed by Flow Cytometry with a low acquisition rate (<500 events/second). Cell cycle status was classified as: G0 (Ki-67^negative^; 2N DNA content), G1 (Ki-67+; 2N DNA content), and S/G2/M (Ki-67+; >2N DNA content). For evaluation of apoptosis, Annexin V staining was performed on surface-stained cells according to manufacturer’s suggestions (Biolegend 640906). BMECs that were DAPI^-^Annexin V^+^ were defined as apoptotic.

### Colony-forming assay

Colony-forming units (CFUs) in semi-solid methylcellulose were quantified to assess hematopoietic progenitor activity. WBM was flushed from femurs and tibiae using a 26.5-gauge needle with MACS buffer. Viable cell counts were determined with a hemocytometer using Trypan Blue (Life Technologies). WBM (5×10^4^ cells) were plated in duplicate wells of Methocult GF M3434 methylcellulose (StemCell Technologies) according to the manufacturer’s suggestions. Colonies were scored for phenotypic CFU-GEMM, CFU-GM, CFU-G, CFU-M, and BFU-E colonies using a SZX16 Stereo-Microscope (Olympus).

### BM transplantation assays

To obtain WBM for transplantation studies, long bones (femur and tibia) were flushed using a 26.5-gauge needle with MACS buffer. For HSC transplantation, WBM was depleted of lineage-committed hematopoietic cells using a Lineage Cell Depletion Kit (Miltenyi), stained with antibodies raised against SCA1 (D7; Biolegend), cKIT (2B8; Biolegend), CD150 (TC15-12F12.2; Biolegend), and CD48 (HM48-1; Biolegend), and HSCs (defined as DAPI^-^lineage^-^SCA1^+^cKIT^+^CD150^+^CD48^-^) were FACS sorted to purity. For primary transplantations, lethally-irradiated (950 Rads) CD45.1 recipients (12-16 weeks) were injected via retro-orbital sinus with either 100 CD45.2^+^ phenotypic HSCs or 5×10^5^ CD45.2^+^ WBM cells along with a 5×10^5^ CD45.1^+^ WBM cell competitive dose. Multi-lineage engraftment was monitored at 4, 8, 16, and 24 weeks post-transplantation by flow cytometry analysis of PB stained with antibodies raised against CD45.2 (104; Biolegend), CD45.1 (A20; Biolegend), and TER119 (TER119; Biolegend) or (CD45.2 (104; Biolegend), GR1 (RB6-8C5; Biolegend), CD11B (M1/70; Biolegend), B220 (RA3-6B2; Biolegend), and CD3 (17A2; Biolegend). For secondary transplantations, WBM cells from primary recipients were isolated and 2×10^6^ total cells per recipient were injected into either wild-type control or *mTOR^(ECKO)^* lethally-irradiated mice (12-16 weeks). Multi-lineage hematopoietic engraftment was monitored at 24 weeks post-transplantation by flow cytometry analysis of peripheral blood as described above.

### Transcriptome Analysis

For HSC RNA isolation, HSCs were FACS sorted to purity as described above, into TRIzol LS Reagent (ThermoFisher Scientific). RNA was purified from TRIzol LS according to the manufacturer’s recommendations. For BMEC RNA isolation, mice were injected retro-orbitally with 25 µg of a fluorophore-conjugated antibody raised again VE-cadherin (BV13; Biolegend) 15 minutes prior to sacrifice. Long bones were isolated, crushed, and enzymatically disassociated in Digestion buffer for 15 minutes at 37°C. Resulting cell suspensions were filtered (40 µm), washed using MACS buffer. Single-cell WBM suspensions were depleted of lineage-committed hematopoietic cells using a Lineage Cell Depletion Kit (Miltenyi) according to the manufacturer’s suggestions. Resulting lineage^-^ cells were surface-stained with fluorophore-conjugated antibodies raised against CD31 (390; Biolegend) and CD45 (30-F11; Biolegend). Stained cells were washed in MACS buffer. BMECs were sorted to purity into TRIzol LS Reagent (ThermoFisher Scientific). RNA was purified from TRIzol LS according to the manufacturer’s recommendations. For gene expression analysis, cDNA was generated and amplified from purified total RNA using the Ovation Pico WTA System V2 (Nugen) according to the manufacturer’s suggested protocol. For microarray analysis, amplified cDNA was further fragmented and biotinylated using the Encore Biotin Module (Nugen) according to the manufacturer’s recommendations. RNA, amplified cDNA, and fragmented cDNA integrity was confirmed using the Agilent RNA 6000 Nano Kit (Agilent) and 2100 Bioanalyzer (Agilent). Resulting DNA was hybridized to GeneChip Mouse Gene 1.0 ST Arrays (Affymetrix), labeled with R-Phycoerythrin-conjugated streptavidin, and scanned using the Affymetrix GeneChip Scanner 3000 7G. All liquid handling was performed using an Affymetrix GeneChip Fluidics Station 450. Microarray quality control and gene expression analysis were done using Affymetrix Expression Console and Transcriptome Analysis Console software. To confirm microarray analysis, Reverse transcription quantitative PCR (RT-qPCR) analysis was performed using 1 ng amplified cDNA template, 1 µM gene-specific primers, and 1X SYBR Green Master Mix (Applied Biosystems) on a ViiA7 real-time PCR instrument (Applied Biosystems). Appropriate no-template controls were included in all experiments. Primers were designed in-house or obtained from the Harvard Primer Bank (Spandidos et al., 2010). Primers are listed in **Table 1**.

**Table 1:**
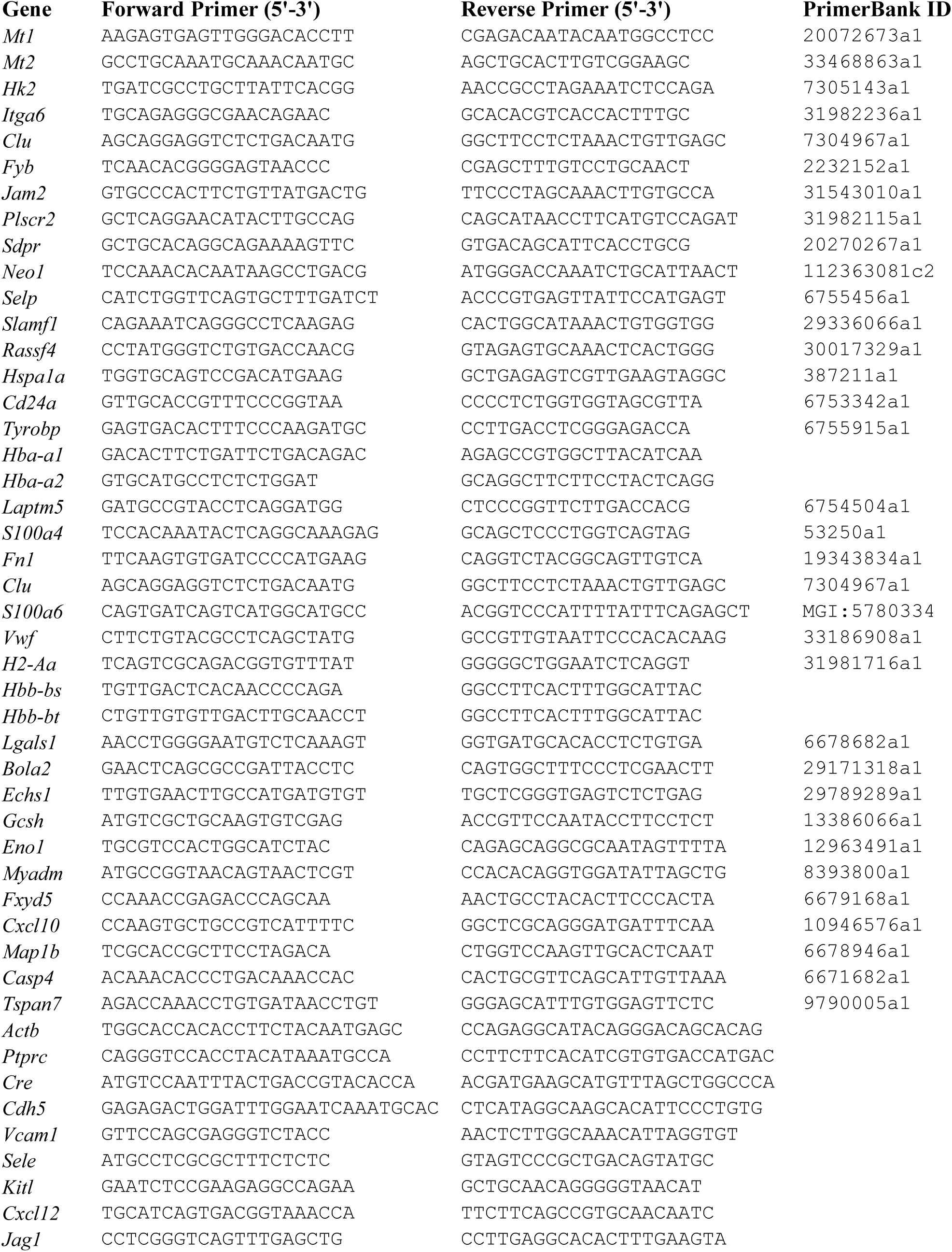
List of Primers.

### Identification of an aged HSC and BMEC gene expression signatures

To define a gene expression signature that characterizes an aged HSC, we compared our microarray expression data with previously published microarray datasets (Chambers et al., 2007; Rossi et al., 2005). Using Venny (http://bioinfogp.cnb.csic.es/tools/venny/), we identified 42 genes showing significant up-regulation and downregulation within aged HSCs that were common between our study and either of these previous studies of which 39 genes showed concordant directional changes across the datasets. 37 out of the 39 genes had detectable mRNA expression in HSCs by RNA-Seq (FPKM >10) (Lis et al., 2017). To define a gene expression signature that characterizes an aged BMEC, we first identified genes that were differentially expressed between young and aged BMECs by utilizing IDEP (Ge et al., 2018). With an FDR cut-off of 0.15 and absolute fold change cut-off of 1.5, we identified 251 genes that were upregulated and 152 genes that were downregulated in aged BMECs, as compared to young BMECs. Reactome Pathway analysis of these differentially expressed genes was performed using WebGestalt (Liao et al., 2019). We compared these differentially expressed genes with a previously published aging meta-analysis (de Magalhaes et al., 2009) utilizing Venny and identified 21 genes showing significant up-regulation in our study and the published dataset, and represent the aged BMEC gene expression signature. We next identified genes that were differentially expressed between young and mTOR^ECKO^ BMECs by utilizing IDEP. With a FDR cut-off of 0.15 and absolute fold change cut-off of 1.5, we identified 74 genes that were upregulated and 29 genes that were downregulated in mTOR^ECKO^ BMECs, as compared to young BMECs. To identify genes that displayed concordant patterns of expression in aged BMECs and mTOR^ECKO^ BMECs, we compared their list of differentially expressed genes using Venny, which demonstrated 14 genes that were upregulated and 5 genes that were down-regulated in both aged BMECs and mTOR^ECKO^ BMECs, as compared to young BMECs. The 14 upregulated genes include *Cd24a, Fn1, Hba-a1, Hba-a2* and *S100a6* that are part of the aged BMEC gene expression signature.

### Immunocytochemistry

To assess γ-H2AX foci and cellular polarity in HSCs, long bones were flushed using a 26.5-gauge needle with MACS buffer. WBM was depleted of lineage-committed hematopoietic cells using a Lineage Cell Depletion Kit (Miltenyi), and then surface-stained with antibodies raised against SCA1 (D7; Biolegend), cKIT (2B8; Biolegend), CD150 (TC15-12F12.2; Biolegend), and CD48 (HM48-1; Biolegend), and HSCs were sorted to purity. A minimum of 1000 sorted HSCs (DAPI^-^lineage^-^ SCA1^+^cKIT^+^CD150^+^CD48^-^) were resuspended in 250 µL of MACS buffer; cytospins were carried out by applying 250 µL resuspended cells to slides using pre-wet Shandon Cytoclip, Sample Chambers, and Filter Cards (ThermoScientific), according to the manufacturer’s suggestions. All cytospins were done using a Shandon Cytospin 4 centrifuge (ThermoScientific) with low acceleration at 1200 rpm for 3 minutes. HSCs were subsequently air dried for 20 minutes on slides, fixed for 10 minutes in 4% paraformaldehyde (PBS, pH 7.2), rinsed three time for 5 minutes in PBS (pH 7.2), and incubated in blocking buffer (10% Normal Goat Serum (Jackson ImmunoResearch) and 0.1% Triton X-100 (Sigma-Aldrich) in PBS (pH 7.2)) for 30 minutes at room temperature. Slides were incubated in primary antibody raised against α-Tubulin (DM1A; Cell Signaling) or phospho-H2AX (Ser139) (JBW301; Millipore) in blocking buffer overnight at 4°C. Cells were washed three times in PBS (pH 7.2) and incubated in blocking buffer for 1 hour with goat anti-mouse IgG (Alexa Fluor 488) (LifeTechnologies), according to the manufacturer’s suggestions. Slides were washed three times in PBS (pH 7.2), stained with DAPI (Biolegend) at 1 µg/mL in PBS (pH 7.2) for 5 minutes at room temperature. Slides were washed three additional times and mounted in ProLong Gold Antifade Mountant (ThermoFisher Scientific). Cells were imaged on a LSM 710 confocal microscope (Zeiss). A minimum of 50 HSCs were analyzed for each genotype. To assess bone marrow vascular morphology, mice were intravenously administered 25 □g of Alexa Fluor 647-conjugated VEcadherin antibody (BV13; Biolegend) via retro-orbital sinus injections. After 15 minutes, mice were euthanized and cardiac perfused with 4% paraformaldehyde (PFA) following which femurs were isolated, stripped of muscle and connective tissue, and fixed in 4% PFA for 30 minutes at room temperature. Bones were washed in 1X PBS 3×5 minutes and cryopreserved in 15% sucrose for 24 hours at 4°C, and further cryopreserved in 30% sucrose for 24 hours at 4°C. Bones were then embedded in a 50% optimal cutting temp (OCT; Tissue-Tek) and 50% sucrose solution and flash frozen in liquid nitrogen. Bones were shaved longitudinally on a Leica CM 3050S cryostat to expose the marrow cavity. Shaved bones were unmounted and washed 2-3 times in 1X PBS until O.C.T. was completely melted. Exposed bones were washed 3×10 minutes in 1X PBS. 40 µm Z-stack whole mount immunofluorescence images of femurs were acquired on a Nikon C2 confocal laser scanning microscope.

### Immunoblot analysis

WBM from long bones (femur and tibia) were flushed using a 26.5-gauge needle with ice-cold PBS (pH 7.2) containing 2 mM EDTA. WBM was depleted of red blood cells (RBC Lysis Buffer; Biolegend) according to the manufacturer’s recommendations. Briefly, flushed marrow cells were pelleted by centrifugation (500g for 5 minutes at 4°C) and the cells were resuspended in 3 mL of ice-cold 1X RBC lysis buffer, vortexed briefly and incubated for 5 minutes on ice. Cells were pelleted by centrifugation (500g for 5 minutes at 4°C), supernatant was discarded, and cells were washed with 3 mL of ice-cold PBS (pH 7.2). Cell pellets were lysed in RIPA buffer (10^7^ cells in 0.5 mL RIPA buffer; Thermo Cat# 89900) containing 2X Phosphatase Inhibitor (Thermo Cat# 78428) and 2X Protease Inhibitor Cocktail (Thermo Cat# 78430) for 1 hour at 4°C with gentle agitation, sonicated, and centrifuged for 10 minutes at 21,000 x g at 4°C to remove insoluble debris. Protein concentrations were determined using the DC Protein Assay (BioRad 5000111) and 20 µg total protein was denatured for 5 min at 70°C in 1X Laemmli Buffer (Sigma Cat # S3401-10VL), resolved on SDS-acrylamide gels and electroblotted to nitrocellulose. Transferred blots were blocked for 1 hour in 5% w/v non-fat dry milk in 1X TBST (Cell Signaling Cat# 9997). Blots were washed 3X for 5 minutes in 1X TBST and incubated overnight at 4°C in 5% BSA w/v in 1X TBST with primary antibodies raised against phospho-S6 (Cell Signaling 4858), S6 (Cell Signaling 2217), phospho 4EBP-1 (Cell Signaling 2855), 4EBP-1 (Cell Signaling 9644) and Actb (Cell Signaling 4970), at the manufacturer recommended dilutions. Blots were washed 3 x 5 mins in 1X TBST and incubated in 5% non-fat dry milk in 1X TBST containing anti-rabbit (H+L) horseradish peroxidase (Jackson ImmunoResearch Laboratories) secondary antibodies at a dilution of 1:20,000 for 1 hour at room temp. Blots were rinsed twice and washed 4 x 5 minutes in 1X TBST and developed using Amersham ECL Prime Western Blotting Detection Reagent (GE Healthcare RPN2232), according to the manufacturer’s suggestions. All blots were developed using Carestream Kodak BioMax Light Film (Sigma-Aldrich).

### Endothelial cell cultures

Primary bone marrow endothelial cell (BMEC) cultures were generated from young (8-12 weeks) C57BL6 mice, as described previously (Poulos et al., 2017). Briefly, femurs and tibiae were gently crushed using a mortar and pestle and digested with Digestion buffer for 15 min at 37°C, filtered (40 µm; Corning 352340), and washed in MACS buffer. WBM was depleted of terminally differentiated hematopoietic cells using a murine Lineage Cell Depletion Kit (Miltenyi Biotec 130-090-858) according to the manufacturer’s recommendations. BM endothelial cells were immunopurified from cell suspensions using sheep anti-rat IgG Dynabeads (ThermoFisher Scientific 11035) pre-captured with a CD31 antibody (MEC13.3; Biolegend) in MACS buffer according to the manufacturer’s suggestions. CD31 selected BM ECs were cultured in endothelial growth media and transduced with 10^4^ pg *myrAkt1* lentivirus per 3×10^4^ ECs. *Akt*-transduced BMECs were selected for seven days in serum- and cytokine-free StemSpan SFEM (StemCell Technologies, Inc. 09650) media. BMEC lines were stained with antibodies against VEcadherin (BV13; Biolegend), CD31 (390; Biolegend), and CD45 (30-F11; Biolegend) and FACS sorted for purity (BMEC defined as CD45-CD31+ VEcadherin+). Established BMEC lines were transduced with GFP-tagged RNAi lentivirus (BLOCK-iT HiPerform Lentiviral Pol II miR RNAi Expression system, Invitrogen Cat# K4934-00) targeting Raptor or Rictor, and transduced cells were selected by FACS sorting for GFP+ cells. DNA oligos for generating RNAi expression vectors were designed using the RNAi designer tool (https://rnaidesigner.thermofisher.com/rnaiexpress/) (Table 2). A panel of 5 lentivirus targeting unique regions within the ORFs of each gene were screened for identifying lentivirus with the highest efficiency of knockdown. As each individual lentivirus demonstrated ∼20-30% knockdown of target genes, vectors were redesigned to co-express two distinct miRNAs from a single lentivirus targeting the same gene by a chaining reaction (Invitrogen Cat# K4938-00). Chained RNAi vectors expressing mRaptor_sh_3 and mRaptor_sh_4 (Raptor) and mRictor_sh_1 and mRictor_sh_4 (Rictor) demonstrated >50% knockdown of the respective target genes in by immunoblotting analysis, and were utilized for experiments. A chained RNAi vector designed to co-express miRNAs targeting the E.Coli *LacZ* gene and a universal negative control sequence (Invitrogen Cat# K4934-00) was utilized to establish ‘control’ BMEC cell lines. Cell lysates were prepared from established BMEC lines utilizing RIPA buffer. Immunoblotting was performed as described above with primary antibodies raised against phospho-Akt (Cell Signaling 9271), Akt (Cell Signaling 4691), phospho 4EBP-1 (Cell Signaling 2855), 4EBP-1 (Cell Signaling 9644), Raptor (Cell Signaling 2280), Rictor (Cell Signaling 2140) and Tubb (Cell Signaling 2146), at the manufacturer recommended dilutions.

**Table 2.**
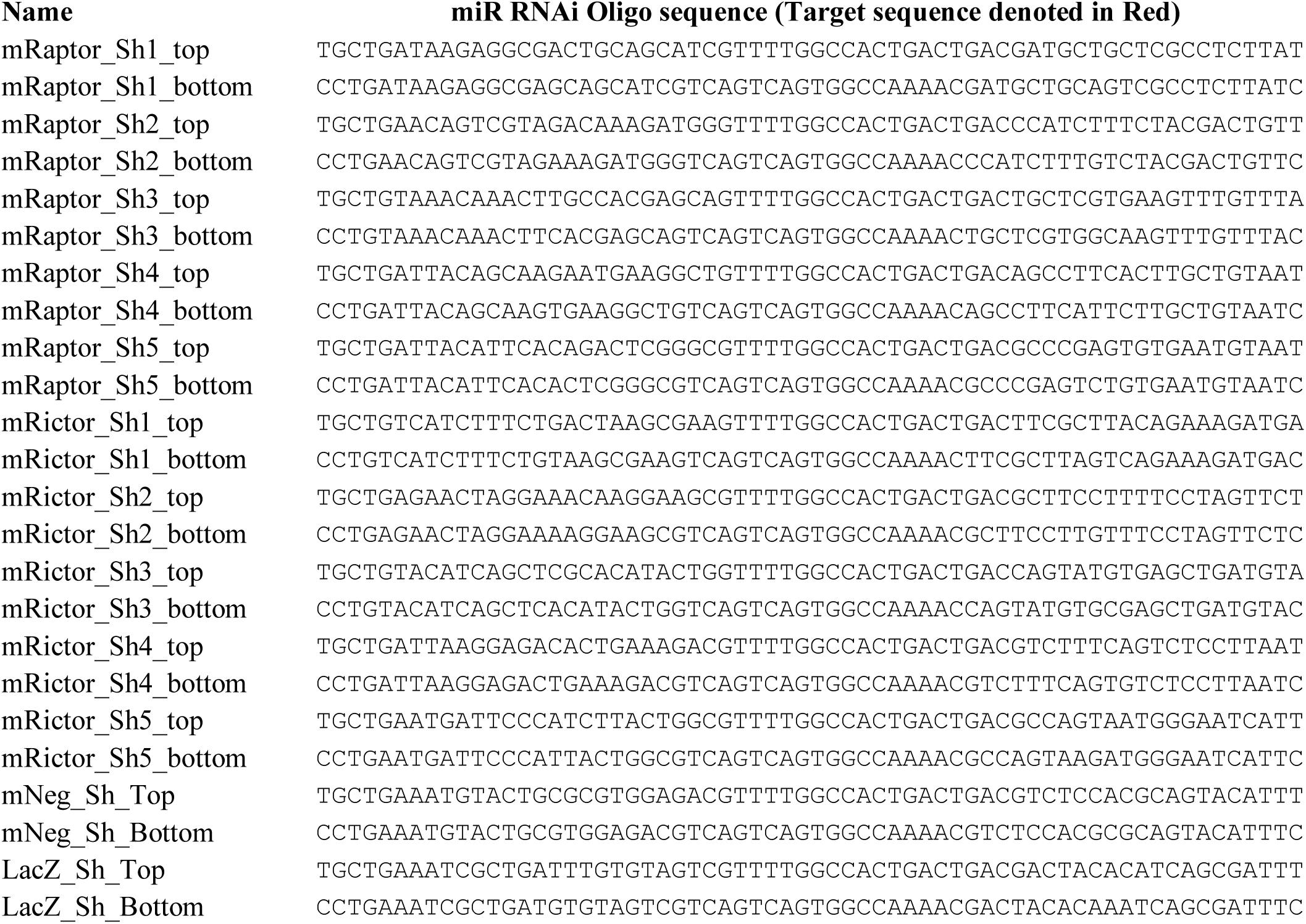
List of DNA oligos synthesized for generating lentiviral miR RNAi expression vectors.

### BMEC-HSPC co-culture

Primary murine HSPCs were co-cultured on established BMEC lines under serum-free conditions with sKITL supplementation, as described previously (Poulos et al., 2016). Briefly, long-bones from young C57BL/6J mice (12 weeks) were isolated and WBM was flushed using a 26.5 gauge needle with MACS buffer. WBM was depleted of lineage+ cells using a Lineage Cell Depletion Kit (Miltenyi Biotec), surface-stained using antibodies raised against cKIT (2B8; Biolegend) and SCA1 (D7; Biolegend), and sorted for KLS populations. FACS-sorted KLS cells (2500 cells/well) were plated on a single well of a 12-well dish pre-plated with BMECs as feeders, in StemSpan SFEM serum-free media (StemCell Technologies) with 50 ng/mL recombinant murine sKITL (PeproTech) and split as needed, as previously described (Poulos et al., 2016). Cells were cultured in 37°C and 5% CO2. All *ex vivo* expansions were analyzed following nine days of co-culture. Total CD45+ cell numbers were calculated after nine days of co-culture, by performing flow cytometry. To assess HSPC activity, total co-culture cells were stained with an antibody raised against CD45.2 (104; Biolegend), and 25,000 FACS-sorted CD45.2+ cells were plated in duplicate wells of Methocult GF M3434 methylcellulose (StemCell Technologies) according to the manufacturer’s suggestions. Colonies were scored for phenotypic CFU-GEMM, CFU-GM, CFU-G, CFU-M, and BFU-E colonies using a SZX16 Stereo-Microscope (Olympus).

### Statistics

Statistical significance was determined using an unpaired two-tailed Student’s t-test, with a significance threshold set at P<0.05. All data are presented as Mean and error bars represent SEM. HSC Microarray gene expression significance was determined using Transcriptome Analysis Console software and a conditioned pair ANOVA test with a significance threshold of P<0.05.

## List of non-standard abbreviations

BMEC: Bone marrow endothelial cells
ECKO: Endothelial cell knockout
GVHD: Graft-versus-host disease
HSCT: Hematopoietic stem cell transplantation
HSPC: Hematopoietic stem and progenitor cells
WBM: Whole bone marrow

## Online Supplemental material

**Figures S1A-D** shows data demonstrating decreased mTOR signaling in the aged BM microenvironment including unfractionated WBM cells, Lineage-CD45+ hematopoietic cells and BM endothelial cells. **Figures S1E-F** shows decreased mTOR signaling within BM cells of aged mice following Rapamycin treatment. **Figures S1G-I** demonstrates that endothelial specific expression of *cre* transgene does not cause hematopoietic defects. **Figures S1J-L** shows that loss of both alleles of endothelial mTOR are essential for inducing HSPC aging phenotypes. **Figure S2** demonstrates that knockdown of either mTORC1 or mTORC2 in cultured endothelial cells, impairs their ability maintain co-cultured HSPCs *ex vivo*. **Figure S3** shows that endothelial specific knockout of mTOR results in delayed hematopoietic recovery and HSPC regeneration following a myelosuppressive injury.

## Author Contributions

M.G.P., P.R., M.C.G.,, L.K., A.G.F., E.L. and J.M.B performed experiments and conducted data analyses. P.R. and J.M.B. conceived the experiments and wrote the manuscript.

## Acknowledgements

Our work is supported by the Tri-Institutional Stem Cell Initiative, American Society of Hematology Scholar Award, American Federation of Aging Scholar Award, National Institutes of Health (1R01CA204308, 1R01HL133021, and 1R01AG065436), and Leukemia and Lymphoma Society. P.R. is supported by Glenn/AFAR scholarship for Research in the Biology of Aging. J.M.B. is a Scholar of the Leukemia and Lymphoma Society. The authors declare no competing financial interests.

## Supplemental Figures and Figure Legends

**Supplemental Figure 1.**
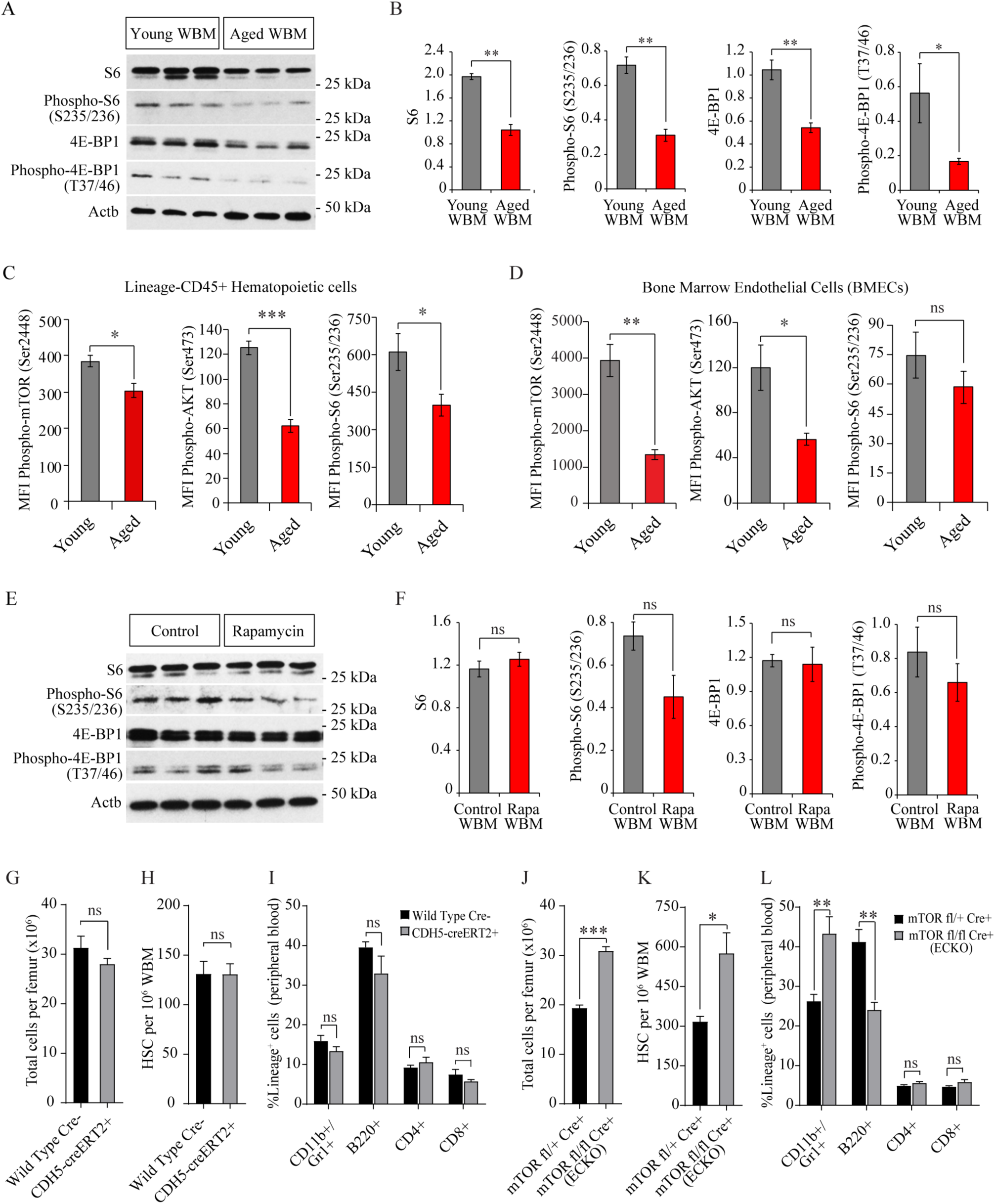
Aging is associated with decreased mTOR signaling within the bone marrow microenvironment. **(A)** Representative immunoblot images demonstrating decreased expression of phospho-S6 and phsopho-4EBP-1 in BM cells of aged mice as compared to young mice. **(B)** Densitometry based quantification of indicated proteins in the BM of aged mice as compared to young mice (n=6 mice per cohort). Expression of *Actb* was used for normalization. Data represents combined analysis of 2 independent experiments. **(C, D)** Quantification of mean fluorescent intensity (MFI) of phospho-mTOR (Ser2448), phospho-AKT (Ser473) and phospho S6 (Ser235/236) by Phospho Flow cytometry in **(C)** Lin-CD45+ HSPCs and **(D)** Lin-CD45-CD31+VECAD+ BMECs (n=5 mice per cohort). **(E)** Representative immunoblot images demonstrating decreased expression of phospho-S6 and phsopho-4EBP-1 in BM cells of aged mice treated with Rapamycin. **(F)** Densitometry based quantification of indicated proteins in the BM of aged mice treated with Rapamycin as compared to aged control mice (n=3 mice per cohort). Expression of *Actb* was used for normalization. **(G-I)** Analysis of wild-type *cre*- (N=3) and *CDH5-creERT2*+ (N=3) mice revealed no significant differences in **(G)** BM cellularity, **(H)** HSC frequency, and **(I)** peripheral blood lineage composition indicating that endothelial-specific expression of *cre-ERT2* transgene does not affect hematopoiesis. **(J-L)** Hematopoietic analysis of heterozygote *mTOR^fl/+^ cre-ERT2+* (N=5) and *mTOR^(ECKO)^* (N=5) demonstrated that unlike *mTOR^(ECKO)^* mice, the littermate heterozygote *mTOR^fl/+^ cre-ERT2+* mice do not manifest increased BM cellularity **(J)**, increased HSC frequency **(K)**, and myeloid-skewed peripheral blood lineage composition **(L)**. All mice utilized in **(G-L)** were administered 200 mg/kg tamoxifen via intraperitoneal injection at a concentration of 30 mg/mL in sunflower oil on three consecutive days, followed by three days off, and three additional days of injection. Note that the same regimen induces HSPC aging phenotypes in homozygote *mTOR^fl/fl^cre-ERT2+* (**Figures 3, 4**), indicating that loss of both alleles of endothelial *mTOR* are essential to induce HSPC aging phenotypes. Error bars represent mean ± SEM. Statistical significance determined using Student’s t-test. *P<0.05, **P<0.01, ***P<0.001, n.s. denotes not statistically significant.

**Supplemental Figure 2.**
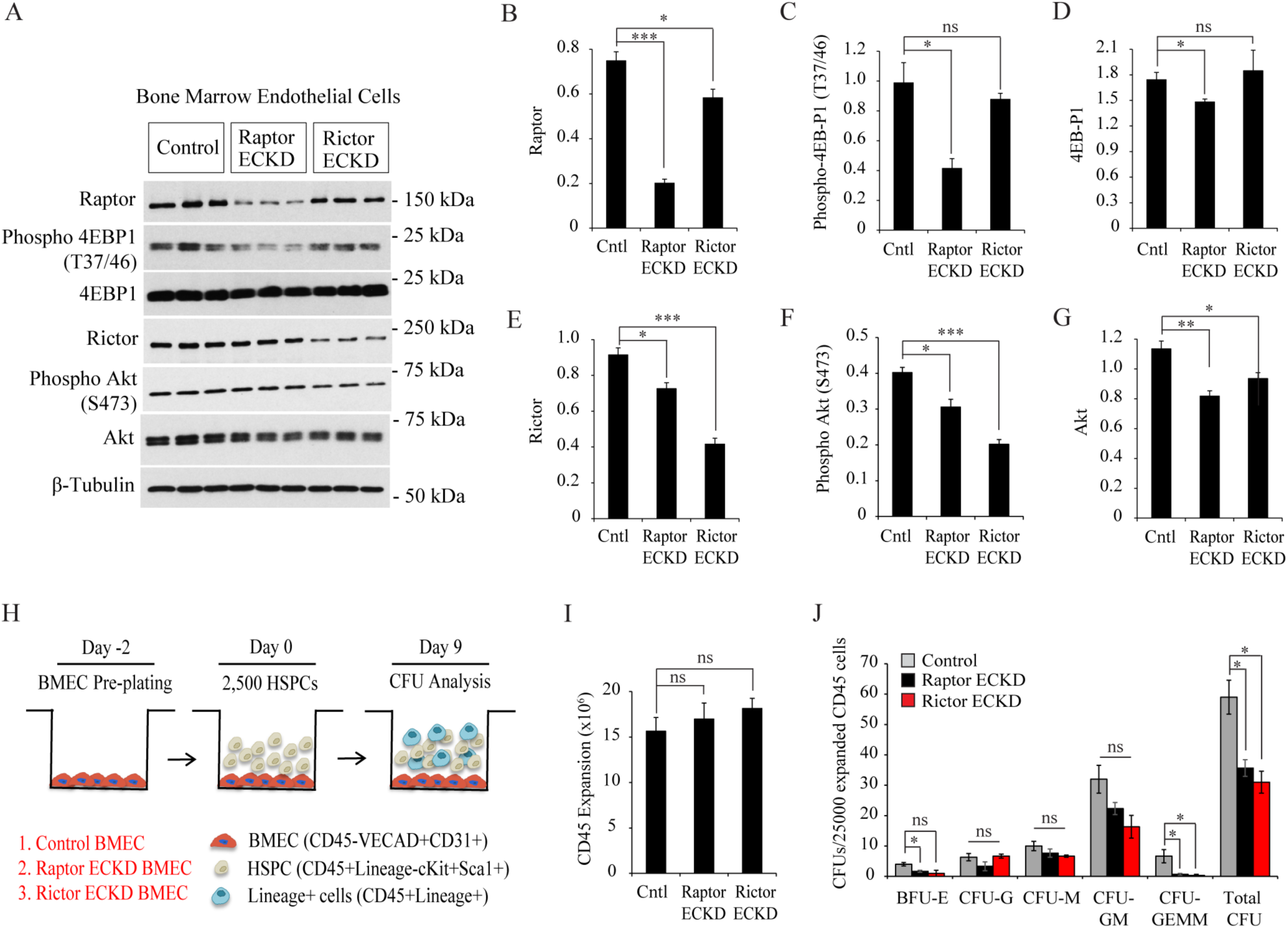
Endothelial mTORC1 and mTORC2 play non-redundant roles in HSPC maintenance. **(A)** Representative immunoblot images demonstrating decreased expression of phospho-4EBP-1 and phospho-Akt in cultured BMECs transduced with lentiviral shRNA targeting Raptor and Rictor, respectively. ECKD denotes ‘Endothelial Cell Knock-Down’.**(B-G)** Densitometry based quantification of indicated proteins. Expression of *Tubb* was used for normalization. **(H)** Schematic of *ex vivo* co-culture assay. In short, HSPCs were co-cultured for 9 days with 50 ng/mL sKitL in serum-free media, followed by phenotypic and functional analysis (n=3 independent co-cultures per genotype). **(I)** Total CD45+ hematopoietic cells estimated by flow cytometry after 9 days of co-culture (n=3 expansions per genotype). **(J)** Day 9 expanded CD45 cells were FACS purified from feeders. 25,000 CD45+ cells were plated in methylcellulose and scored for CFUs after 8 days (n=3 per genotype). Data in **A-J** represent combined analysis of 3 independent experiments. Error bars represent sample mean ± SEM. Statistical significance determined using Student’s t-test. * P<0.05; ** P<0.01; *** P<0.001, n.s. denotes not statistically significant.

**Supplemental Figure 3.**
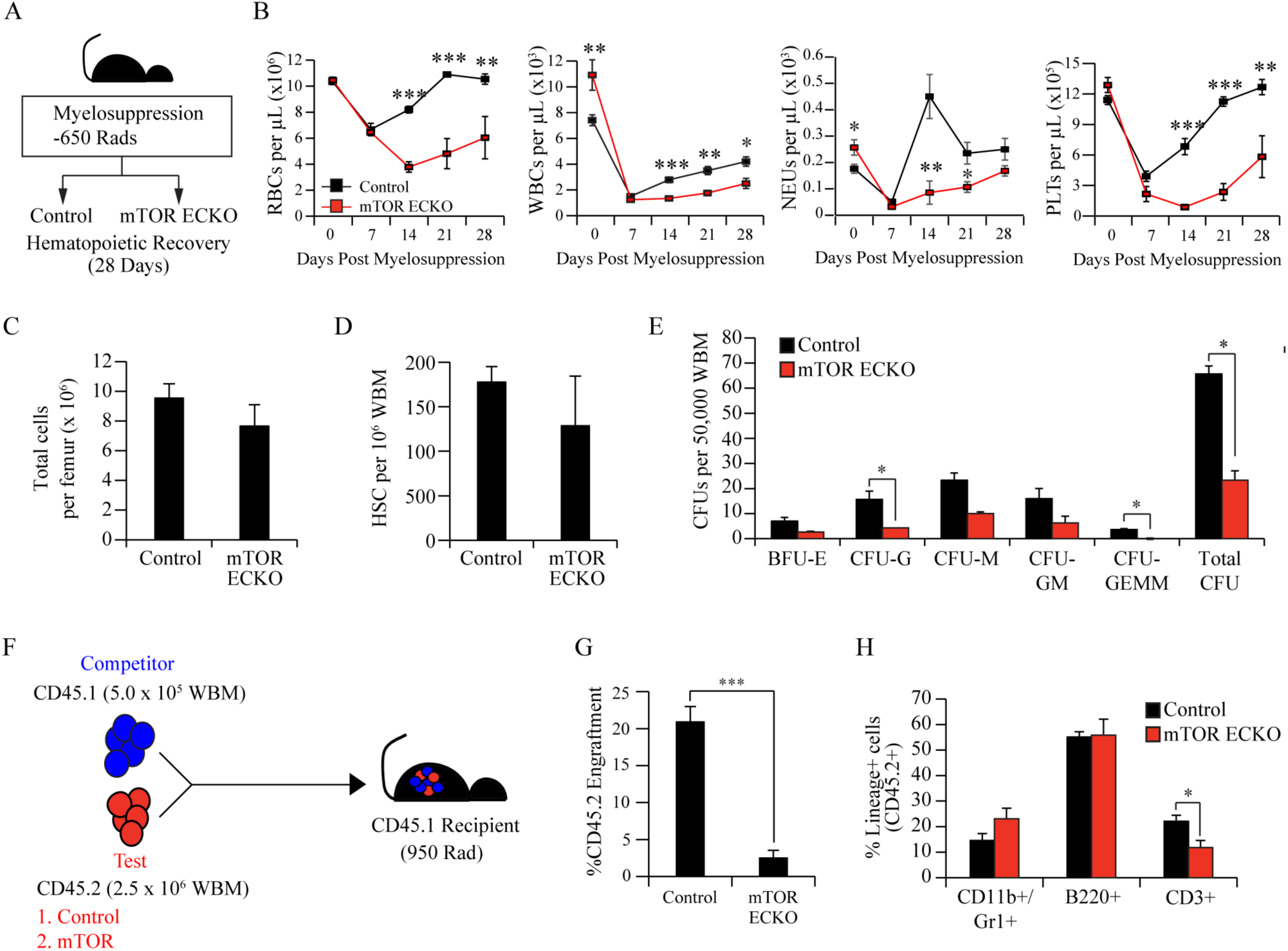
Loss of endothelial mTOR adversely impacts hematopoietic recovery following myelosuppressive injury. **(A)** Experimental design to assess hematopoietic recovery following radiation induced myelosuppressive injury. **(B)** Peripheral blood counts demonstrate a significant delay in red blood cell, white blood cell and platelet recovery in young *mTOR^(ECKO)^* mice following radiation (n=5 mice per cohort). **(C)** Total hematopoietic cells per femur and **(D)** the frequency of HSCs per femur (n=5 per cohort) after 28 days following myelosuppressive injury. **(E)** Methylcellulose-based colony assay to assess hematopoietic progenitor activity (n=3 mice per cohort). Data in **B-E** represent combined analysis of 2 independent experiments. **(F)** Schematic of competitive WBM transplantation assay to determine long-term engraftment potential and multi-lineage reconstitution ability of BM cells following myelosuppressive injury. **(G-H)** Peripheral blood analysis after 16 weeks post-transplantation revealed a loss of long-term engraftment potential **(G),** along with impaired reconstitution of CD3+ cells **(H),** in donor BM cells derived from *mTOR^(ECKO)^* mice (n=10 recipients per cohort). Data represents average engraftment following transplantation of cells derived from n=5 independent donor mice per cohort. Error bars represent sample mean ± SEM. Statistical significance determined using Student’s t-test. * P<0.05; ** P<0.01; *** P<0.001.

